# Disruption of the *Pseudomonas aeruginosa* Tat system perturbs PQS-dependent quorum sensing and biofilm maturation through loss of the Rieske cytochrome *bc_1_* sub-unit

**DOI:** 10.1101/2021.03.01.433341

**Authors:** Eliza Ye-Chen Soh, Frances Smith, Maxime Rémi Gimenez, Liang Yang, Rebecca Munk Vejborg, Matthew Fletcher, Nigel Halliday, Sophie Bleves, Stephan Heeb, Miguel Camara, Michael Givskov, Kim R. Hardie, Tim Tolker-Nielsen, Bérengère Ize, Paul Williams

## Abstract

Extracellular DNA (eDNA) is a major constituent of the extracellular matrix of *Pseudomonas aeruginosa* biofilms and its release is regulated via the pseudomonas quinolone signal (PQS) dependent quorum sensing (QS). By screening a *P. aeruginosa* transposon library to identify factors required for DNA release, mutants with insertions in the twin-arginine translocation (Tat) pathway were identified as exhibiting reduced eDNA release, and defective biofilm architecture with enhanced susceptibility to tobramycin. *P. aeruginosa tat* mutants showed substantial reductions in pyocyanin, rhamnolipid and membrane vesicle (MV) production consistent with perturbation of 2-heptyl-3-hydroxy-4-quinolone (PQS) dependent QS as demonstrated by changes in *pqsA* expression and 2-alkyl-4-quinolone (AQ) production. Provision of exogenous PQS to the *tat* mutants did not return *pqsA*, *rhlA* or *phzA1* expression or pyocyanin production to wild type levels. However, transformation of the *tat* mutants with the AQ-independent *pqs* effector *pqsE* restored *phzA1* expression and pyocyanin production. Since mutation or inhibition of Tat prevented PQS-driven auto-induction, we sought to identify the Tat secretion substrate responsible. A *pqsA::lux* fusion was introduced into each of 34 validated *P. aeruginosa* Tat substrate deletion mutants. Analysis of each mutant for reduced bioluminescence revealed that the signalling defect was associated with the Rieske iron-sulfur subunit of the cytochrome *bc_1_* complex. In common with the parent strain, a Rieske mutant exhibited defective PQS signalling, AQ production, *rhlA* expression and eDNA release that could be restored by genetic complementation. Thus, lack of the Rieske sub-unit export is clearly responsible for the Tat-mediated perturbation of PQS-dependent QS, the loss of virulence factor production, biofilm eDNA and the tobramycin tolerance of *P. aeruginosa* biofilms.

**Author Summary:** *Pseudomonas aeruginosa* is a highly adaptable human pathogen responsible for causing chronic biofilm-associated infections. Biofilms are highly refractory to host defences and antibiotics and thus difficult to eradicate. The biofilm extracellular matrix incorporates extracellular DNA (eDNA). This stabilizes biofilm architecture and helps confer tolerance to antibiotics. Since mechanisms that control eDNA release are not well understood, we screened a *P. aeruginosa* mutant bank for strains with defects in eDNA release and discovered a role for the twin-arginine translocation (Tat) pathway that exports folded proteins across the cytoplasmic membrane. Perturbation of the Tat pathway resulted in defective biofilms susceptible to antibiotic treatment as a consequence of perturbed pseudomonas quinolone (PQS) signalling. This resulted in the failure to produce or release biofilm components including eDNA, phenazines and rhamnolipids as well as microvesicles. Furthermore, we discovered that perturbation of PQS signalling was a consequence of the inability of *tat* mutants to translocate the Rieske subunit of the cytochrome *bc_1_* complex involved in electron transfer and energy transduction. Given the importance of PQS signalling and the Tat system to virulence and biofilm maturation in *P. aeruginosa*, our findings underline the potential of the Tat system as a drug target for novel antimicrobial agents.

## Introduction

*Pseudomonas aeruginosa* is an opportunistic pathogen that causes a wide range of human infections including lung, urinary tract and wound, bacteremia and infections associated with medical devices [1]. It is notorious for its tolerance to antimicrobial agents, a property that is largely a consequence of its ability to form biofilm communities [1, 2]. Bacterial exoproducts including cell surface appendages, extracellular polymeric substances, biosurfactants and secondary metabolites all contribute to *P. aeruginosa* biofilm formation and maturation [3–7].

Apart from exopolysaccharides such as Psl, Pel and alginate, the extracellular polymeric matrix of *P. aeruginosa* biofilms incorporates proteins, rhamnolipids, membrane vesicles (MVs) and extracellular DNA (eDNA) [5,8–10]. Rhamnolipid biosurfactants are required during the initial stages of micro-colony formation and are involved in the migration-dependent formation of the caps of mushroom-shaped micro-colonies formed in flow-cell grown biofilms [10]. They also aid the maintenance of channels between multicellular structures within biofilms and contribute to biofilm dispersal [5]. With respect to the biofilm micro-colonies that characteristically form in flow-chambers fed with glucose minimal medium, eDNA is present at high concentrations in the outer layers of microcolonies in young biofilms. However, in mature biofilms, eDNA is primarily located in the stalks at the borders between micro-colony caps and stalks [8].

The release of eDNA occurs via the lysis of a sub-population of bacterial cells [10–14]. It is involved in attachment, aggregation and stabilization of biofilm microcolonies. eDNA can act as a nutrient source, chelate metal cations and confer tolerance to antibiotics such as the polymyxins and aminoglycosides [10,12,13]. eDNA also binds other biopolymers (exopolysaccharides and proteins) stabilizing biofilm architecture and conferring protection against adverse chemical and physical challenges [12, 13]. By intercalating with eDNA, secondary metabolites such as phenazines enhance biofilm integrity [12, 15]. Pyocyanin for example can contribute to DNA release through the formation of reactive oxygen species such as hydrogen peroxide that damage cell membranes [12]. Although the mechanism(s) responsible for eDNA release has not been fully elucidated both eDNA and MVs can be generated via explosive cell lysis mediated via a cryptic prophage endolysin encoded within the R- and F-pyocin gene clusters [14].

In *P. aeruginosa*, rhamnolipids and pyocyanin production, eDNA and MV release, and hence biofilm development, are all controlled by quorum sensing (QS) [1,8,16]. Consequently, *P. aeruginosa* mutants with defects in this cell-to-cell communication system form aberrant, flat undifferentiated biofilms [10]. In *P. aeruginosa*, the QS regulatory network consists of a hierarchical cascade incorporating the overlapping *las*, *rhl* and *pqs* pathways that employ *N*-acylhomoserine lactones (AHLs) and 2-alkyl-4-quinolones (AQs) as signal molecules [1,16,17]. All three QS systems contain auto-induction loops whereby activation of a dedicated transcriptional regulator by the cognate QS signal molecule induces expression of the target synthase such that QS signal molecule production can be rapidly amplified to promote co-ordination of gene expression at the population level.

*P. aeruginosa* produces a diverse family of AQs and AQ *N*-oxides [18, 19] of which 2-heptyl-3-hydroxy-4-quinolone (the *Pseudomonas* Quinolone Signal, PQS) and its immediate precursor 2-heptyl-4-hydroxyquinoline (HHQ) are most closely associated with PQS signalling [17]. Most of the genes required for AQ biosynthesis (*pqsABCDE*) and response (*pqsR;* also called *mvfR*) are located at the same genetic locus although *pqsH* and *pqsL* are distally located [17]. PqsA catalyses the formation of anthraniloyl-CoA that is condensed with malonyl-CoA by PqsD to form 2-aminobenzoylacetyl-CoA (2-ABA-CoA) [20, 21]. The latter is converted to 2-aminobenzoylacetate (2-ABA) via the thioesterase functionality of PqsE [22]. The PqsBC heterodimer condenses 2-ABA with octanoyl-coA to generate HHQ [23, 24]. PQS is formed through the oxidation of HHQ by PqsH [25] while formation of the AQ *N*-oxides such as 2-heptyl-4-hydroxyquinoline *N*-oxide (HQNO) requires the alternative mono-oxygenase PqsL [18]. The PqsE protein has dual functions; while it is not essential for AQ biosynthesis, it is required for the AQ-independent production of several factors that contribute to biofilm maturation including pyocyanin, rhamnolipids and lectin A [26]. While activation of RhlR-dependent genes depends on PqsE [27], the AQ-independent, thioesterase-independent mechanism by which PqsE acts has not yet been elucidated [17, 22].

The *pqs* system is subject to positive autoinduction, since the LysR-type transcriptional regulator PqsR (MvfR), binds to the promoter region of *pqsABCDE* (P*pqsA*) triggering transcription once activated by HHQ or PQS [28–30] (**Fig. 1**). Therefore, by analogy with other QS systems, HHQ and PQS can both act as autoinducers by generating a positive feedback loop that accelerates their biosynthesis. However, in contrast to HHQ which only regulates the *pqsABCDE* operon [17], PQS is a ferric iron chelator [31] that not only drives AQ biosynthesis via PqsR but also the expression of genes involved in the iron-starvation response and virulence factor production *via* PqsR-dependent and PqsR-independent pathways [17]. In addition, PQS can act as a cell-sensitizing pro-oxidant [32] and is required for MV production via a direct physical interaction with lipopolysaccharide (LPS) within the outer membrane [33]. The packaging of PQS within MVs also provides a means for trafficking this hydrophobic QS signal within a *P. aeruginosa* population [34].

**Fig 1.**
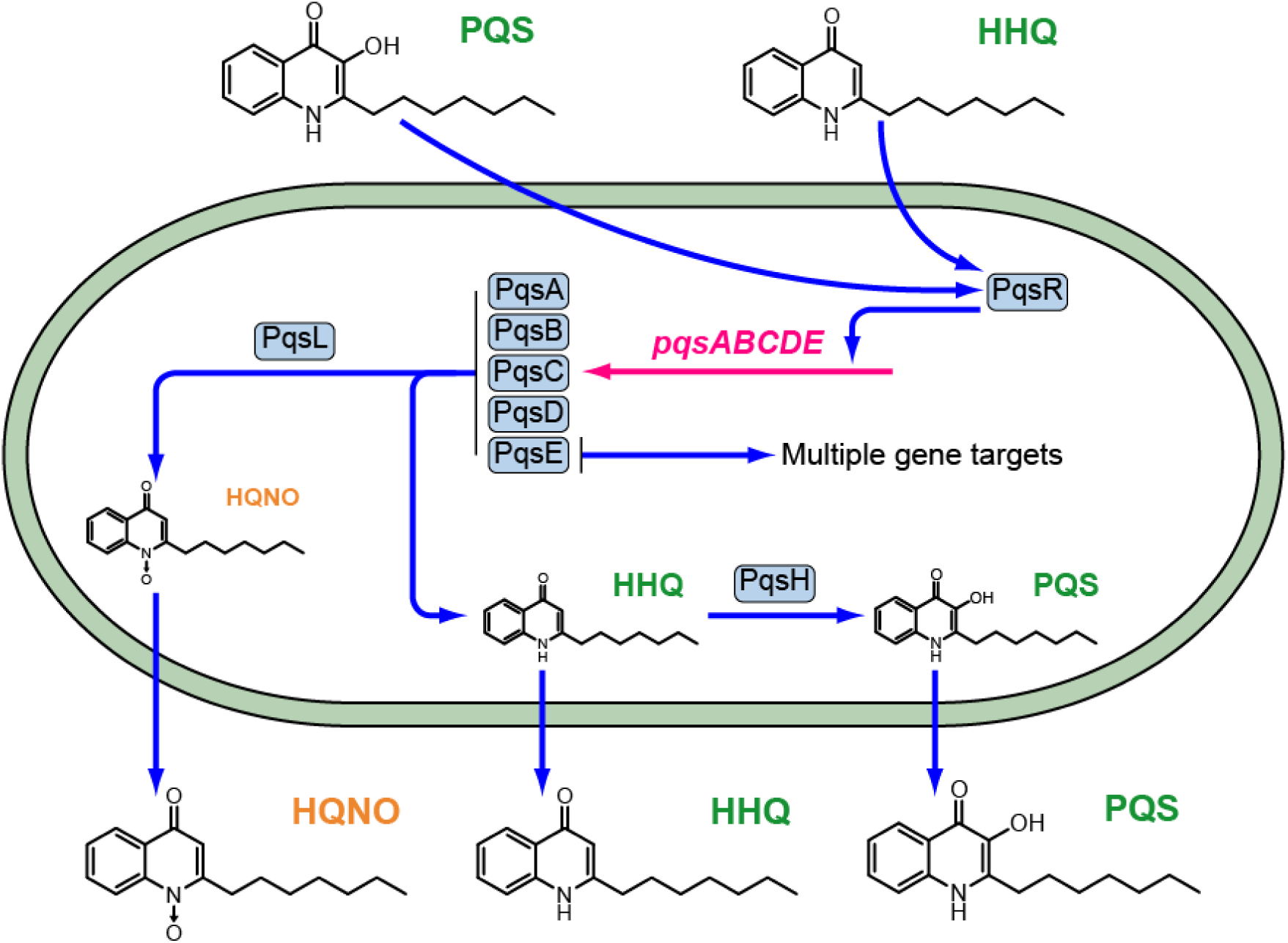
The PQS signalling pathway in *P. aeruginosa*. The PqsABCDE proteins synthesize HHQ, which is converted to PQS by PqsH and also HQNO in conjunction with PqsL. Both HHQ and PQS are released by the cells into the extracellular environment and are taken back up by neighboring cells. Autoinduction occurs when either HHQ or PQS binds to PqsR and amplifies expression of the *pqsABCDE* operon. The *pqsE* gene product has dual functions contributing to AQ biosynthesis as a thioesterase and also via an AQ-independent, thioesterase-independent mechanism to e.g. pyocyanin, rhamnolipid, and lectin production and to biofilm maturation. The conversion of HHQ to PQS confers additional functionalities since PQS unlike HHQ induces microvesicle formation and is a potent iron chelator.

With respect to biofilm development, the *pqs* system is of particular interest because *pqs* biosynthetic mutants fail to produce eDNA, rhamnolipids, pyocyanin and MVs, and form thin defective biofilms containing little eDNA [8, 27,33,35]. The mechanism involved in PQS-mediated DNA-release in biofilms is not understood but has been suggested to be linked to phage induction causing cell lysis [8,36–40]. Although explosive cell lysis releases eDNA in biofilms and generates MVs through vesicularization of shattered membrane fragments, *pqsA* mutants are not defective for explosive lysis [14] and therefore this phenomenon is unlikely to account for PQS-dependent eDNA release.

In the present study we sought to identify additional factors involved in eDNA release by screening a transposon (Tn) mutant library for eDNA-release defective mutants. Apart from *pqs* biosynthetic mutants, we obtained Tn insertion mutants within the twin-arginine translocation (Tat) pathway that exhibited reduced levels of eDNA release, fail to produce rhamnolipids or pyocyanin and form defective, eDNA- poor, antibiotic susceptible biofilms. Since mutation or deletion of *tat* resulted in altered AQ production, reduced pyocyanin, rhamnolipid and MVs, and as the *tat* mutants were refractory to exogenously supplied PQS, the aberrant biofilm phenotype observed could be accounted for by perturbation of PQS autoinduction. By screening a library of *P. aeruginosa* Tat substrate mutants, we identified the Rieske sub-unit of the cytochrome *bc_1_* complex as being required for *pqs*-dependent QS and hence eDNA release and biofilm maturation.

## Results

### Transposon mutagenesis screen for P. aeruginosa mutants exhibiting reduced DNA release

To identify *P. aeruginosa* genes that contribute to eDNA release, a mariner transposon (Tn) mutant library was generated in strain PAO1. Approximately 10,000 mutants grown in microtitre plates were assayed for reduced eDNA release using propidium iodide (PI) to quantify eDNA because it is unable to penetrate live bacteria and because its fluorescence is enhanced 30-fold on binding DNA [41]. From the initial screen, 84 Tn insertion mutants were selected and re-screened to eliminate strains with double Tn insertions and to confirm their eDNA phenotype. For 34 of the remaining mutants, the regions flanking each Tn insertion were sequenced and the corresponding genes identified. For many of the eDNA-release deficient mutants, the Tn insertions were located within genes required for AQ biosynthesis (*pqsC* and *pqsH)* or regulation (*pqsR)* (**Fig. 2**) These data confirm our previous work that first uncovered a role for PQS signalling in eDNA release [8]. Apart from the *pqs* mutants, two mutants were obtained with insertions in the *tatA* and *tatB* genes respectively (**Fig. 2**). These code for components of the twin-arginine translocation (Tat) system that mediates the export of folded proteins and was originally named with respect to the presence of an Arg-Arg motif in the signal sequence of Tat-exported products also called Tat substrates [42, 43]. In *P. aeruginosa* diverse proteins involved in phosphate and iron metabolism, virulence and energy transduction are exported to the periplasm, or secreted via the Tat export system and *tat* mutants exhibit pleiotropic phenotypes [44, 45]. **Fig. 2** shows that genetic complementation of the *P. aeruginosa tatA* mutant with a plasmid-borne copy restored eDNA release.

**Fig 2.**
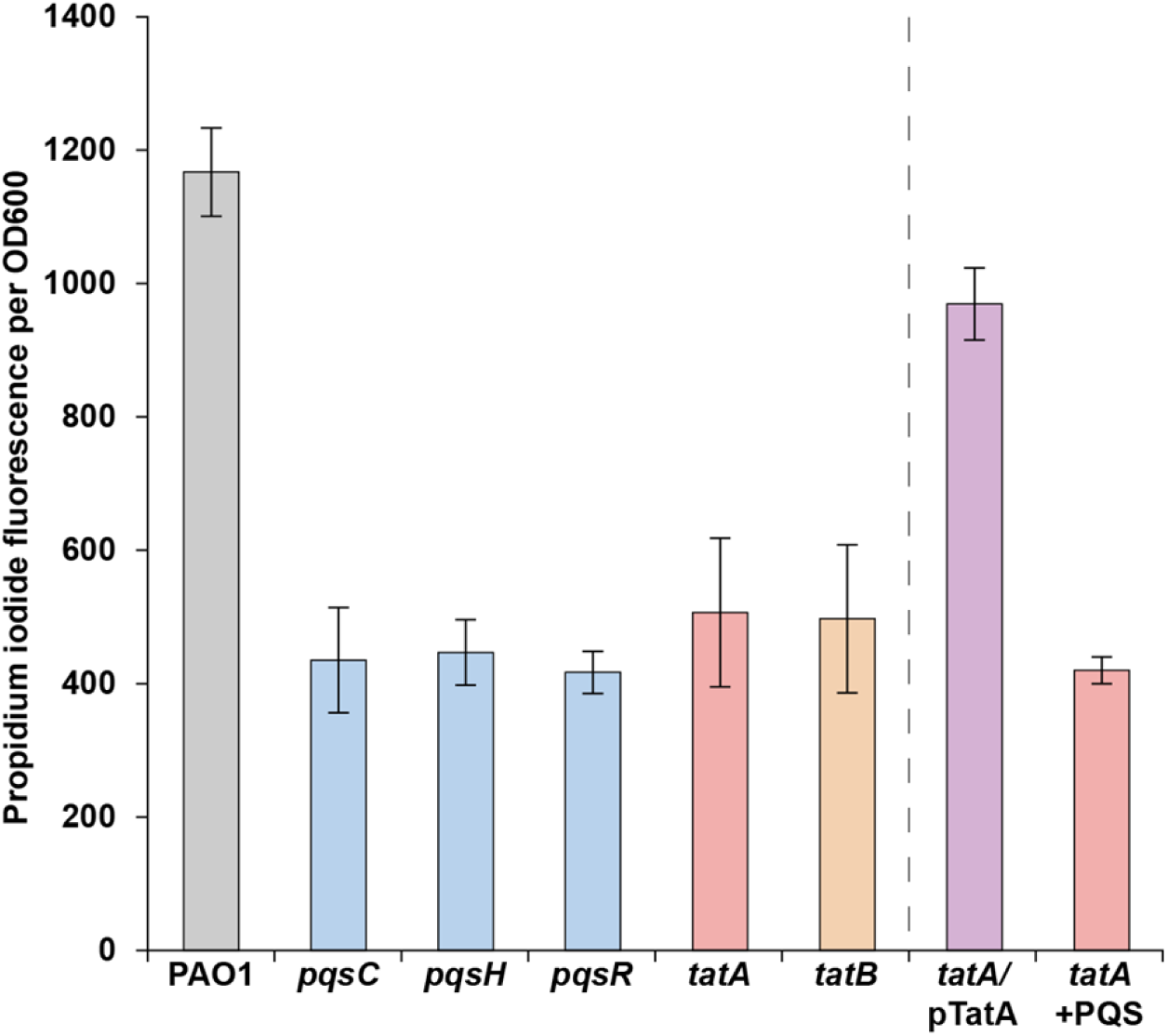
Transposon mutant screen for *P. aeruginosa* strains defective for eDNA release. *P. aeruginosa* wild-type and mutant strains were grown for 24 h in 96 well microtiter plates containing ABTG medium, after which the relative levels of eDNA in the cultures were determined using a PI binding assay. The means and standard deviations of eight replicates are shown.

### The Tat pathway contributes to biofilm development in P. aeruginosa

Since eDNA makes an important contribution to biofilm development and architecture [10,12,13], biofilm formation by the *P. aeruginosa tatA* mutant in flow-chambers was investigated. After 4-days growth, the *P. aeruginosa* wild-type and complemented *tatA*/pTatA mutant formed biofilms with mushroom-shaped structures whereas the *tatA* mutant formed thin, flat biofilms (**Fig. 3A**). Ethidium bromide staining of the latter revealed that they contained much less eDNA than the structured biofilms formed by the wild-type and complemented *tatA* mutant strain (**Fig. 3A**). Consistent with this biofilm phenotype, exposure of the biofilms formed by each of the three strains to tobramycin showed that the *tatA* mutant biofilm was more sensitive to tobramycin than either the wild-type or *tatA*/pTatA mutant biofilm (**Fig. 3B**).

**Fig 3.**
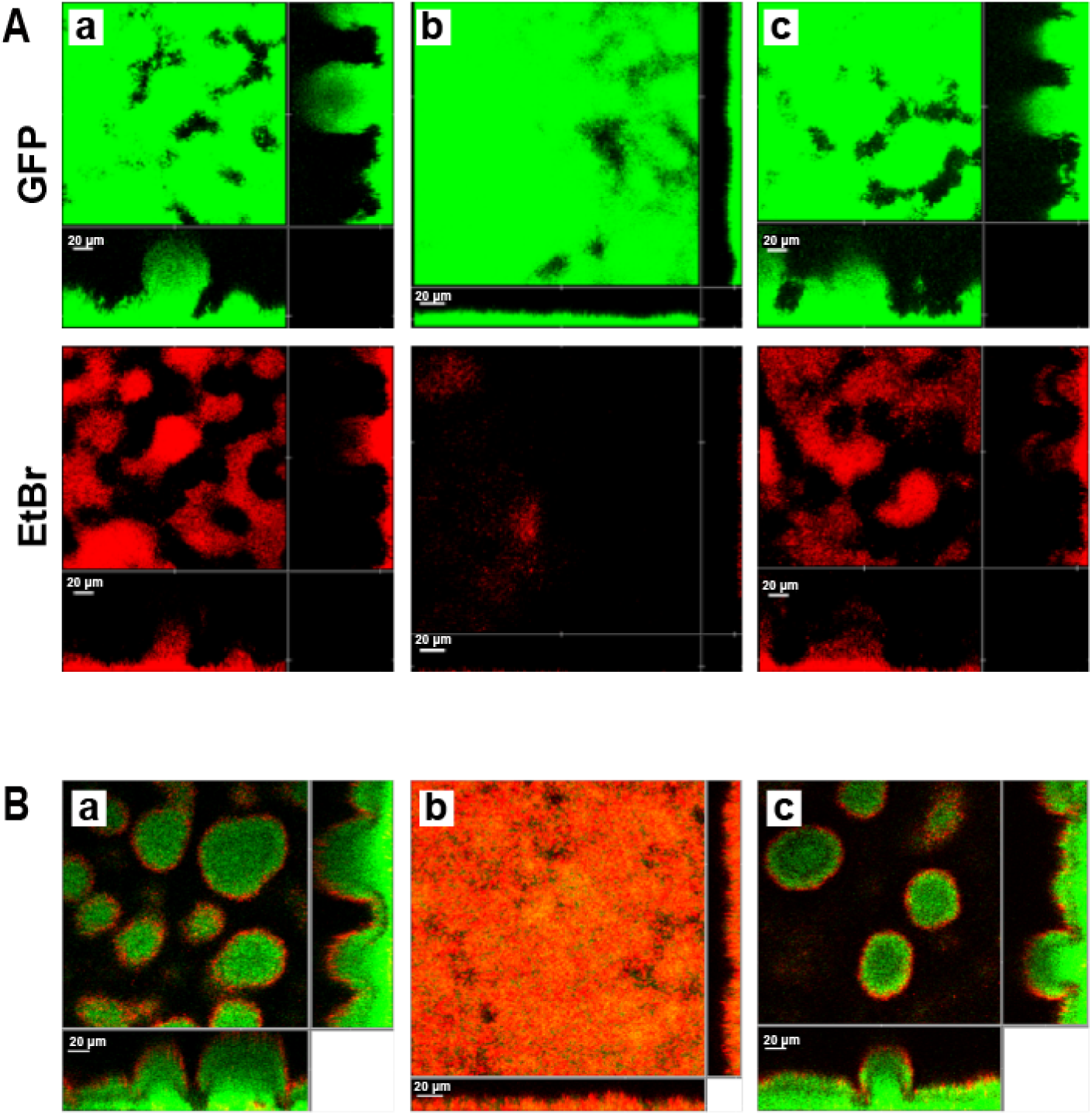
*P. aeruginosa tat* mutants form defective biofilms with increased susceptibility to tobramycin. CLSM images showing four-day-old biofilms formed in flow chambers of the *gfp*-tagged *P. aeruginosa* wild-type (a), *tatA* mutant (b) and genetically complemented *tatA* mutant (c). In **(A)** biofilms were stained for total biomass with Syto9 (green) and for eDNA with ethidium bromide (red). In **(B)** biofilms were treated with tobramycin and the medium was supplemented with propidium iodide prior to CLSM such that dead cells appear red while live cells appear green. Each panel shows one horizontal optical section two flanking vertical optical sections. Bars, 20 µm.

### P. aeruginosa tatA mutants are defective in the production of rhamnolipids, pyocyanin and MVs

Since rhamnolipids, pyocyanin and MVs are all important components of *P. aeruginosa* biofilms, their production was quantified in the *tatA* Tn insertion mutant and in a Δ*tatABC* deletion mutant. **Fig. 4 (A and B)** shows that both *tat* mutants produce substantially less pyocyanin and rhamnolipid than the parent or *tatA* complemented strain. Furthermore, MV levels (**Fig. 4C**) were reduced by ∼50% in the *tatA* mutant compared with the wild type and could be restored by genetic complementation.

**Fig 4.**
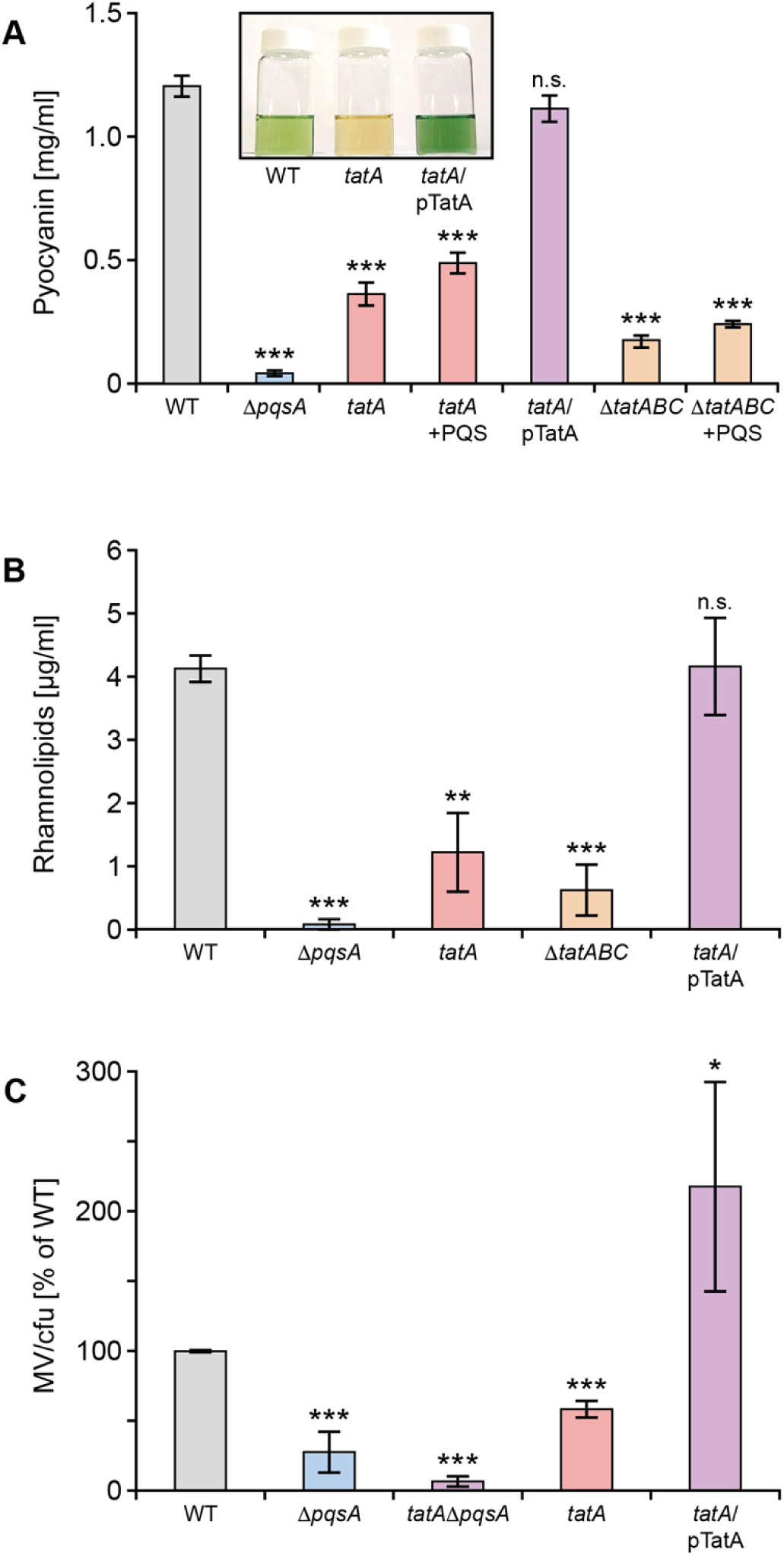
Production of pyocyanin (**A**), rhamnolipids (**B**) and MVs (**C**) are reduced in *P. aeruginosa tat* mutants. (**A**) Pyocyanin levels are shown in the *P. aeruginosa* wild type, Δ*pqsA*, *tatA*, and *tatABC* deletion mutants and the *tatA* mutant complemented with plasmid-borne *tatA*. The impact of exogenous PQS (40 μM) on the *tatA* and Δ*tatABC* mutants is also shown. Insert panel shows the absence of green pigment in the *tatA* mutant compared with the wild type and complemented *tatA* mutant. (**B**) Rhamnolipid production in the *tatA* and Δ*tatABC* mutants compared with the wild type, Δ*pqsA* mutant and *tatA* complemented with plasmid-borne *tatA.* (**C**) Comparison of MV production in the *tatA* mutant, complemented *tatA* mutant and in a double *tatA* Δ*pqsA* mutant compared with the wild type strain. Experiments were repeated in triplicate at least twice. ***p < 0.001, **p < 0.01; n.s. not significant.

### Inactivation of the tat pathway by mutation or small molecule-mediated inhibition perturbs PQS signalling

The reductions in eDNA release, rhamnolipids, pyocyanin and MVs noted in the *tat* mutant as well as its biofilm phenotype are similar to those observed for *P. aeruginosa* strains with mutations in *pqs* genes such as *pqsA*, the first gene in the AQ biosynthetic pathway (see **Figs. 2** and **Fig 4** and [8, 40]). These data suggested that the *tat* mutant biofilm phenotype was likely, at least in part, to be due to a defect in PQS signalling.

To investigate the impact of the *tatA* mutation on the expression of *pqsA,* a CTX::*pqsA’-lux* fusion was introduced into the chromosomal CTX site of both the wild type and *tatA* mutant. **Fig. 5A** shows that *pqsA* expression in the *tatA* mutant is reduced ∼4 fold compared with the wild type strain and restored by genetic complementation of the mutant. In agreement with these data, the Tat inhibitor, Bayer 11-7082, identified by Vasil *et al* [46] reduced *pqsA* expression in the wild type PAO1 strain by ∼4 fold consistent with the reduction noted for the CTX::*pqsA-lux* fusion in the *tatA* mutant (**Fig. 5B**). Bayer-11 7082 had no effect on growth or light output in *P. aeruginosa* expressing the *lux* genes from a derepressed *lac* promoter (**S1 Fig.**). In addition, the concentrations of PQS in whole culture extracts of *P. aeruginosa* after growth in LB medium as determined by LC-MS/MS was respectively ∼56% lower in the *tatA* mutant compared with the wild type and complemented *tat* mutant (**Fig. 5C**). Since *pqsA* expression and hence AQ production is also PqsR/MvfR-dependent, we compared the expression of *pqsR* in the Δ*tatABC* with the parent strain and found no difference (**S2 Fig.**).

**Fig 5.**
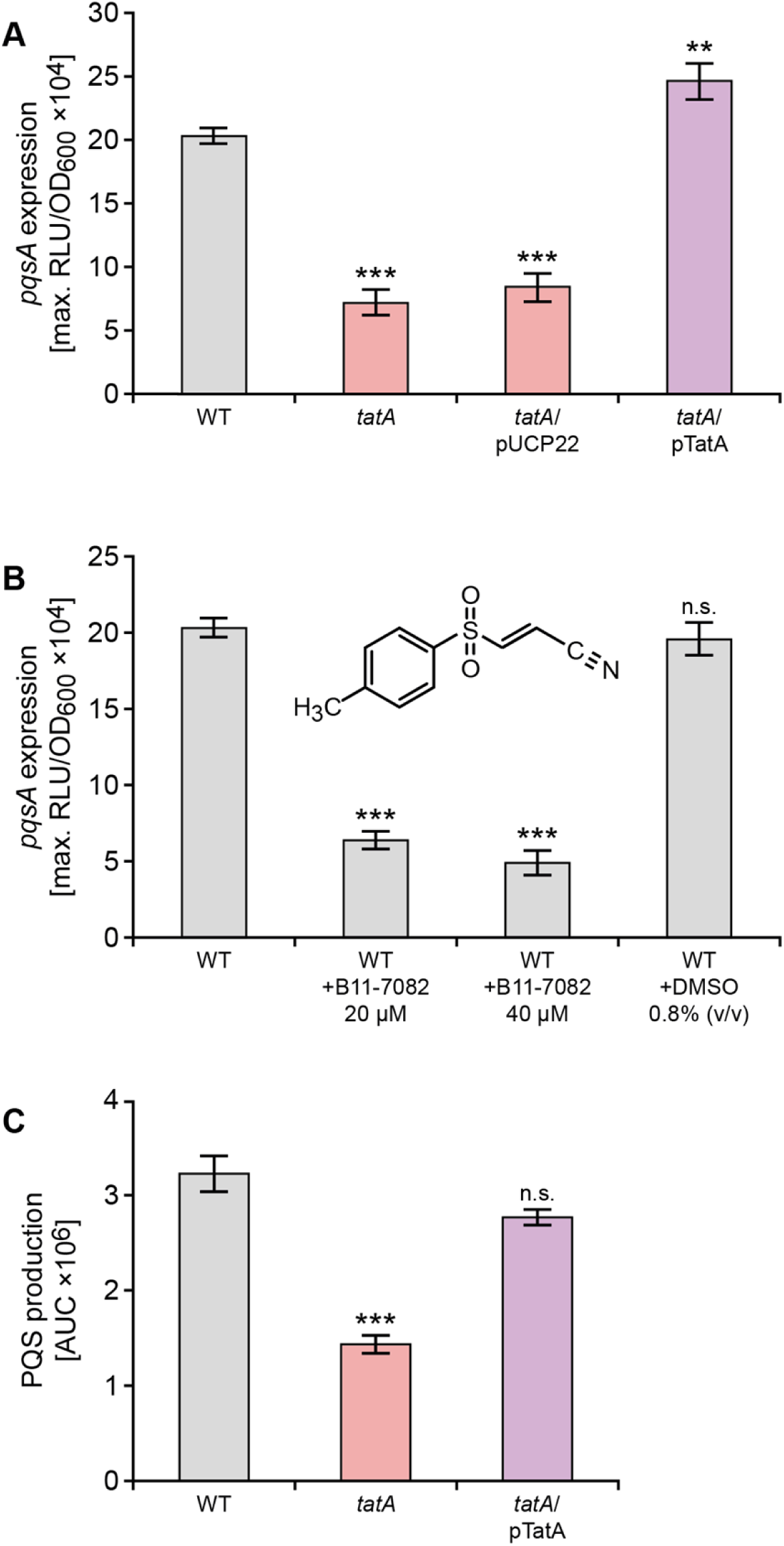
Mutation of *tatA* or exposure to the Tat inhibitor Bayer 11-7082 inhibits *pqsA* expression and AQ production. (**A**) Mutation of *tatA* or (**B**) treatment with Bayer 11-7082 supplied at either 20 μM or 40 μM reduces the maximal expression of a *P. aeruginosa* PAO1 chromosomal *pqsA’-lux* promoter fusion without affecting growth. (**C**) LC-MS/MS analysis of PQS production by *P. aeruginosa* PAO1 wild type compared with the *tatA* mutant and complemented *tatA* mutant. Experiments were repeated in triplicate at least twice. ***p < 0.001; n.s. not significant.

Since mutation of *tatA* results in reduced *pqsA* expression, it is possible that the auto-induction of PQS signalling via PqsR/MvfR is compromised. To uncouple the autoinduction of AQ production, the *pqsABCD* genes were introduced into the *P. aeruginosa* Δ*pqsA* and Δ*pqsA*Δ*tatA* mutants respectively on a plasmid (pBBRMCS5::*pqsABCD*) that constitutively expresses the *pqsABCD* genes [47]. **S3 Fig.** shows that PQS, HHQ and HQNO are present in the culture medium of the Δ*pqsA* mutant transformed with pBBRMCS5::*pqsABCD*. However neither the cell free supernatant nor whole cells of the *tatA* Δ*pqsA* double mutant transformed with pBBRMCS5::*pqsABCD* contained or accumulated intracellular AQs (**S4 Fig.**) suggesting that in the absence of autoinduction in the *tat* mutant background. *P. aeruginosa* is unable to synthesize AQs.

### Exogenous PQS does not restore PQS signalling in a P. aeruginosa *tatA* mutant

QS systems are characteristically autoinducible such that exogenous provision of the cognate signal molecule induces expression of the signal synthase and hence downstream target genes [48]. When the *tatA* mutant was provided with exogenous PQS, eDNA release did not increase (**Fig. 2**). To investigate this finding further, either PQS or HHQ was exogenously supplied to the wild type, *tatA* mutant or the complemented *tatA* mutant strains carrying chromosomal *pqsA’-lux* fusions. The data presented in **Fig. 6A** show that the response of the *tatA* mutant to PQS or HHQ respectively at 5 or 20 μM with respect to *pqsA* expression was at least 2-fold lower than the controls. Since both wild type and *tatA* mutant still produce AQs endogenously, the experiments were repeated in the *P. aeruginosa* Δ*pqsA* and *tatA* Δ*pqsA* mutants since no AQs are produced in these genetic backgrounds. **Fig. 6B** shows that the response to both PQS and HHQ is substantially reduced (e.g. ∼4 fold at 5 μM PQS) in the absence of *tatA* in the *P. aeruginosa tatA* Δ*pqsA* double mutant.

**Fig 6.**
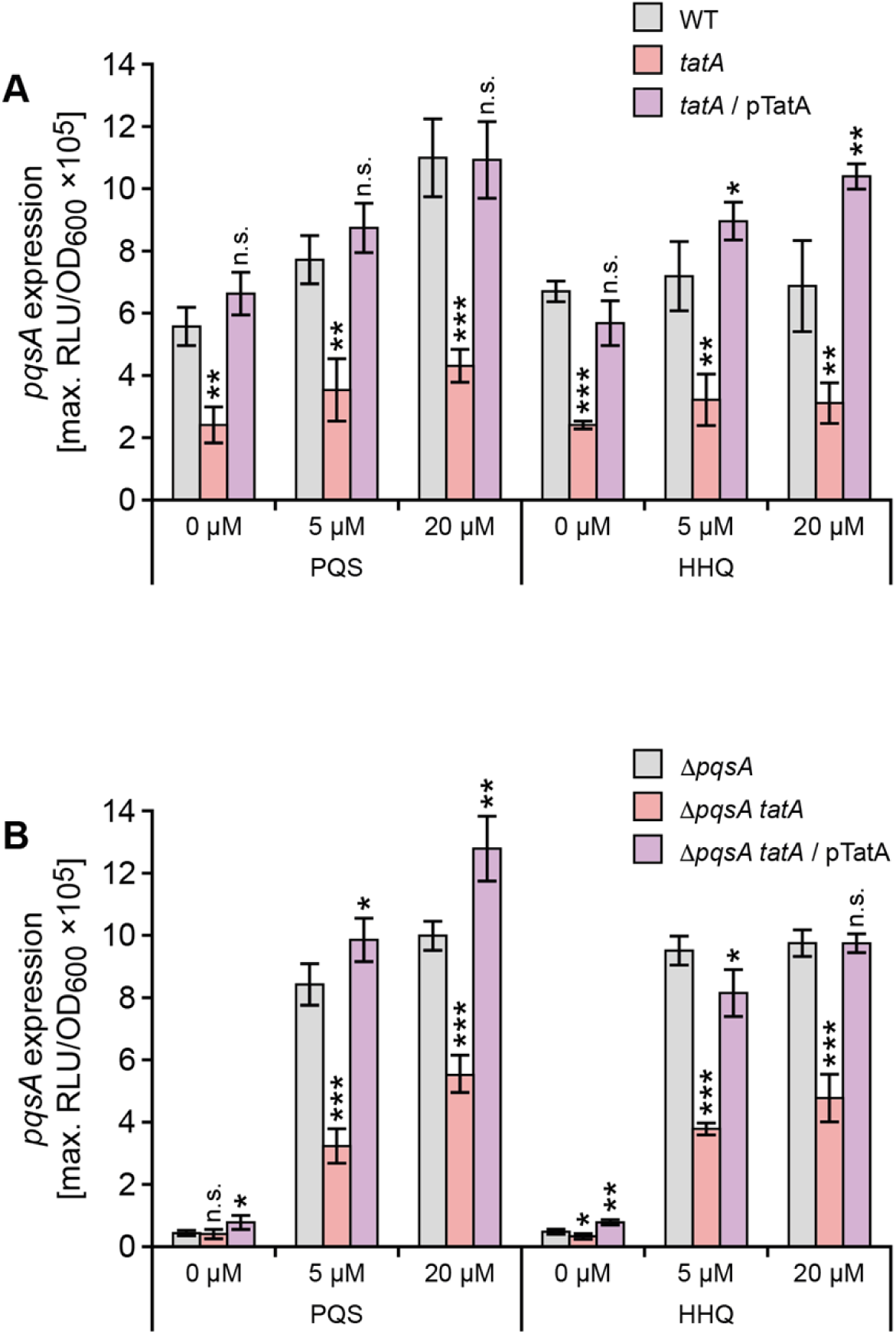
Exogenous AQs do not restore *pqsA* expression in *P. aeruginosa tatA* (A) or *tatA* Δ*pqsA* (B) mutants. Exogenous PQS or HHQ was added at 5 μM or 20 μM to **(A)** wild type, the *tatA* mutant and complemented *tatA* mutant or (**B**) Δ*pqsA*, Δ*pqsA tatA* or *tatA* Δ*pqsA* mutant complemented with *tatA.* Maximal light output from the chromosomal *pqsA’-lux* fusion was recorded as a function of growth (RLU/OD_600_). Experiments were repeated in triplicate at least twice. ***p < 0.001, **p < 0.01, and *p < 0.05; n.s. not significant.

To determine the consequences of perturbed PQS signalling on the expression of the rhamnolipid (*rhlA*) and pyocyanin biosynthetic genes (*P. aeruginosa* has two, almost identical redundant 7 gene phenazine biosynthetic operons termed *phzA1-G1* and *phzA2-G2;* [49]), the corresponding miniCTX-*lux* promoter fusions for *rhlA* and *phzA1* respectively were constructed and introduced onto the chromosomes of the wild type, Δ*pqsA* and *tatA* Δ*pqsA* mutants respectively. **Fig. 7A** shows the expression profiles of *rhlA’-lux* as a function of time. Both the wild type and Δ*pqsA* mutant show an ∼2 fold increase in *rhlA* expression when supplied with exogenous PQS (20 μM) and share similar profiles over the growth curve. In contrast, the *rhlA’-lux* fusion in the *tatA* mutant does not show the same expression profile or response to exogenous PQS as the wild type and Δ*pqsA* mutant strains. The *rhlA’-lux* expression profile in the *tatA* mutant supplied with exogenous PQS is however clearly restored when the mutation is complemented by pTatA (**Fig. 7A**).

**Fig 7.**
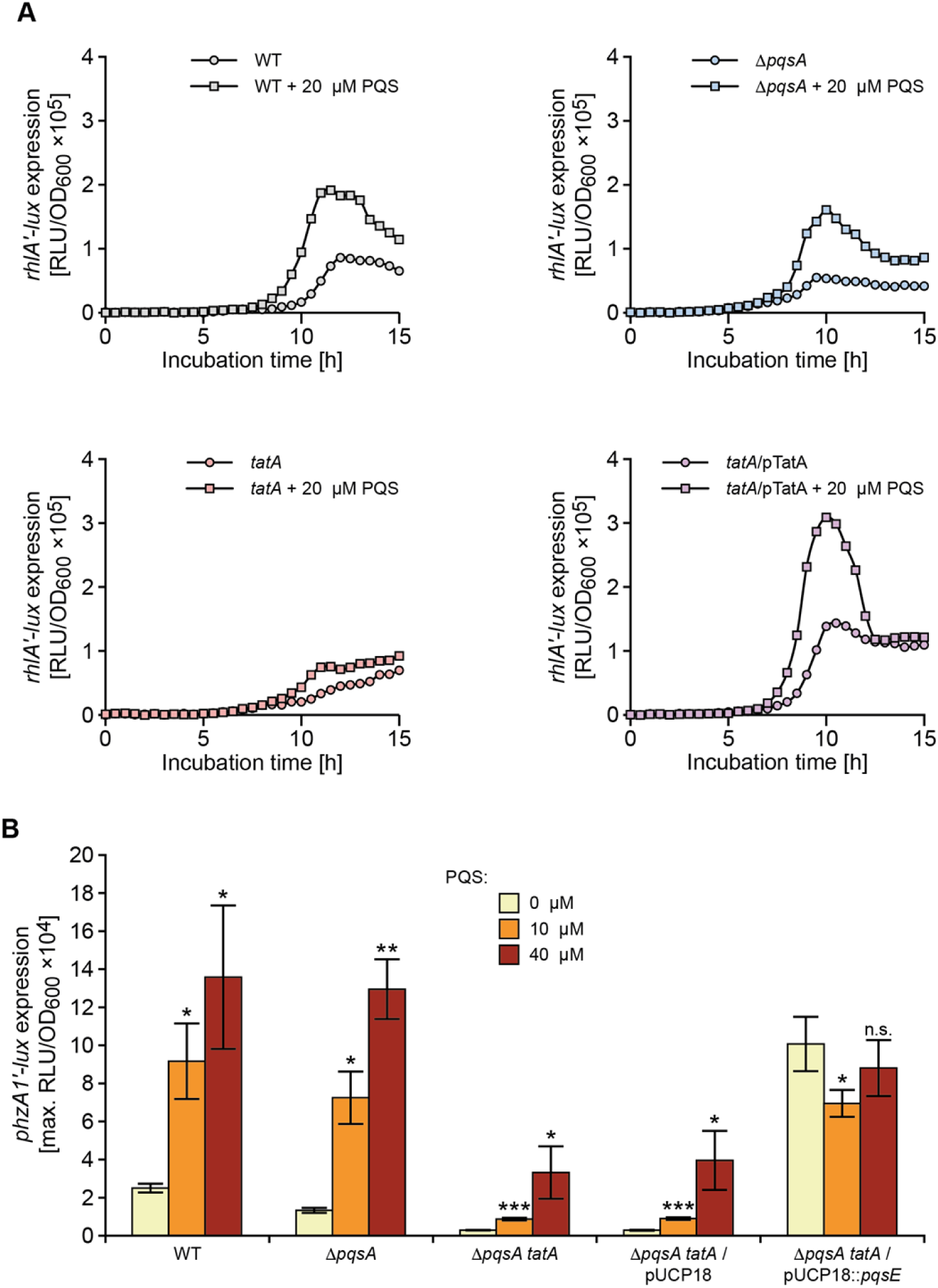
Rhamnolipid (*rhlA*) and pyocyanin biosynthetic (*phzA1*) genes show altered expression profiles in *P. aeruginosa tat* mutants and fail to respond to exogenous PQS. (**A**) Light output from a chromosomal *rhlA’-lux* fusion as a function of growth (RLU/OD) over time when introduced into the wild type, *pqsA* and *tatA* mutants or complemented *tatA* mutant in the absence or presence of exogenous PQS (20 μM). **(B)** Maximal light output from a chromosomal *phzA1’-lux* fusion as a function of growth (RLU/OD_600_) when introduced into the wild type, *pqsA*, or *pqsA tatA* mutants in the absence or presence of exogenous PQS (10 or 40 μM) or plasmid-borne *pqsE* or the pUCP18 vector control. Experiments were repeated in triplicate at least twice. ***p < 0.001, **p < 0.01, and *p < 0.05; n.s. not significant.

AQ-dependent QS is required for *phzA1* expression [27, 49]. Exogenous PQS increased *phzA1’-lux* expression by ∼4 fold in both wild type and Δ*pqsA* mutant backgrounds (**Fig. 7B**). However, the *tatA* Δ*pqsA* double mutant responded comparatively poorly to PQS (**Fig. 7B**).

### PqsE restores pyocyanin in the tat mutants

Although PqsE is not essential for AQ biosynthesis, it is required for the production of pyocyanin, rhamnolipids and biofilm maturation and its function is independent of PQS, HHQ or PqsR [27, 50]. Consequently, the expression of certain PQS signalling-dependent exoproducts such as pyocyanin can be restored in a *pqsA* negative (and hence AQ-negative) mutant by expressing *pqsE*. To determine whether expression of *pqsE* alone could restore *phzA1* expression and pyocyanin production in the *tat* mutants, *pqsE* was introduced on pUCP18. **Fig. 7B** shows that *phzA1* expression can be restored in the *tatA* Δ*pqsA* mutant in the absence of AQs while **Fig. 8** confirms that plasmid-borne *pqsE* restores pyocyanin production.

**Fig 8.**
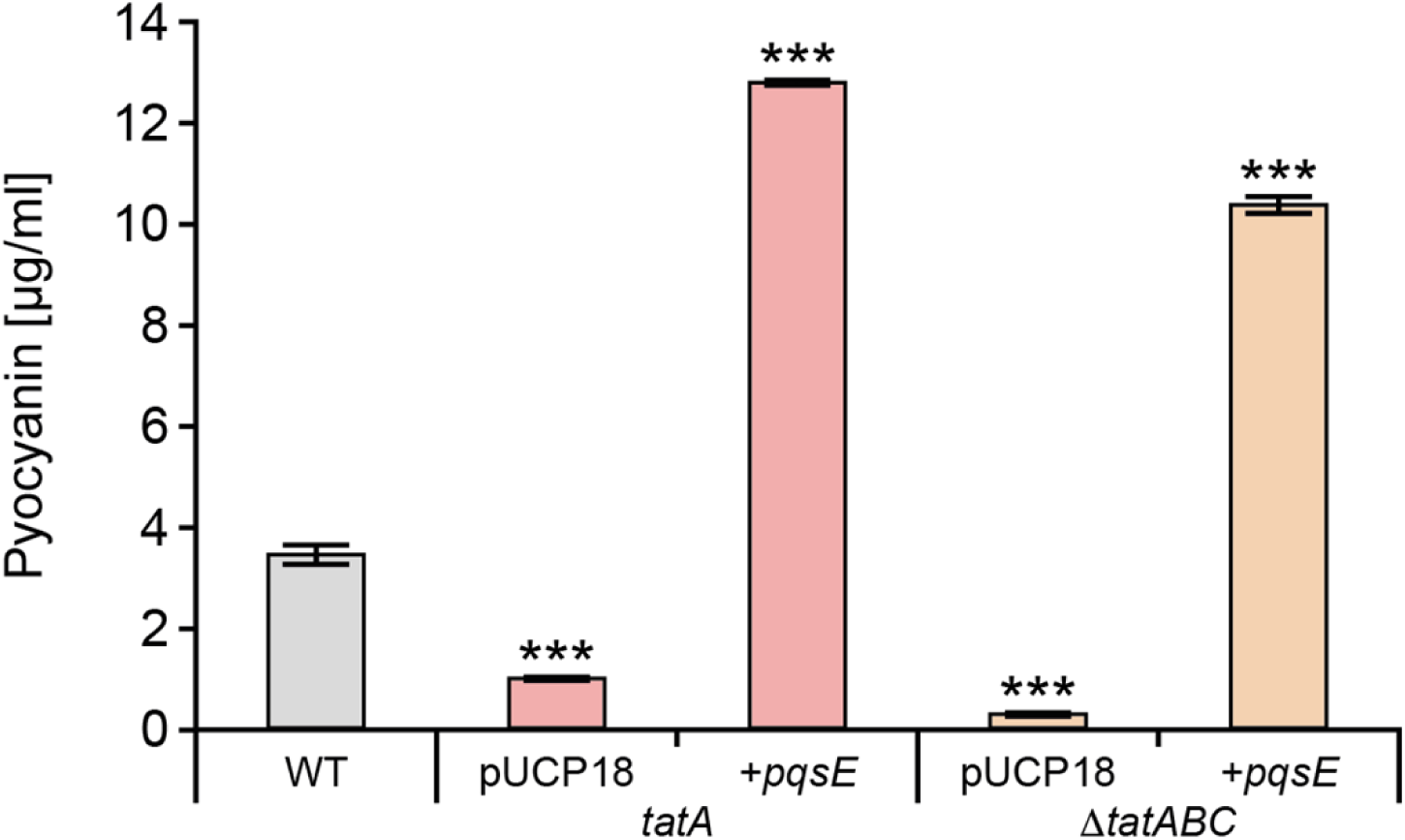
PqsE restores pyocyanin production in *P. aeruginosa tat* mutants. Production of pyocyanin by the wild type compared with the *tatA* and Δ*tatABC* mutants transformed with plasmid-borne *pqsE* or the empty vector. Experiments were repeated in triplicate at least twice. ***p < 0.001,

### The reduction in rhamnolipid production in the *tat* mutants does not account for the perturbation of *PQS signalling*

In *P. aeruginosa* biofilms, rhamnolipids provide protective shielding against neutrophils [51, 52] and contribute to the effectiveness of PQS signalling by enhancing the solubility and bioactivity of PQS [53]. In **Fig. 4B**, we show that rhamnolipid production is substantially reduced in the *P. aeruginosa tat* mutant backgrounds. To determine whether the perturbation of PQS signalling in the *tat* mutants is a consequence of reduced rhamnolipid production, we investigated the impact of exogenous rhamnolipids on *pqsA* expression. **S5 Fig.** shows that the addition of purified rhamnolipids (10 or 50 μg/ml) to the *tatA* Δ*pqsA* mutant with or without PQS (40 μM) had little effect on *pqsA* expression.

### Identification of the Tat substrate responsible for perturbation of *PQS signalling*

Recently Gimenez et al [45] experimentally validated the Tat-mediated export of 34 *P. aeruginosa* gene products predicted to have Tat signal peptides. To determine which of the exported Tat substrates was responsible for perturbation of PQS signalling, allelic replacement mutants were constructed in *P. aeruginosa* strain PA14 for each substrate. Before introducing the *pqsA’-lux* fusion onto the chromosomal CTX site of each Tat substrate mutant, we first confirmed that PQS signalling in a PA14 Δ*tatABC* mutant was perturbed in a similar manner to that observed for PAO1, the genetic background used so far in the study (**Fig. 9A**). Determination of the maximum expression of *pqsA’-lux* for each of the 34 Tat substrate mutants (**Fig. 9A**) revealed that although a number of mutants exhibited reduced light output, the greatest reduction was observed for deletion of PA14_57570 (equivalent to PA4431 in PAO1, here designated *petA* following the nomenclature of orthologues described in other species). This gene codes an iron-sulfur cluster protein, the Rieske subunit of the cytochrome *bc_1_* complex.

**Fig 9.**
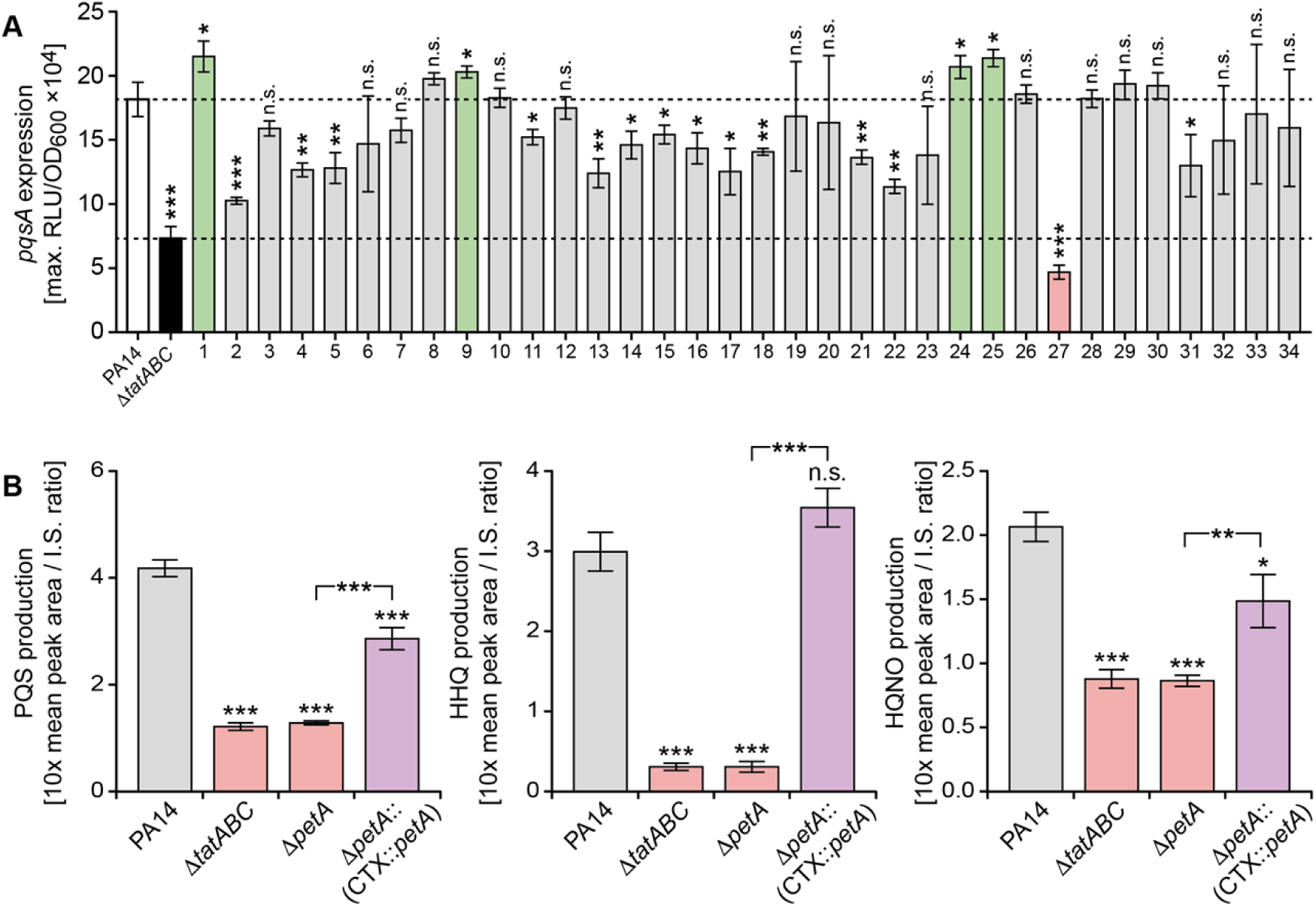
Tat substrate screen for *P. aeruginosa* PA14 mutants with defects in *pqs* signalling uncovers a role for the cytochrome *bc_1_* Rieske sub-unit. (**A**) Comparison of peak *pqsA* expression in each of 34 Tat substrate mutants (**Table S3**) transformed with a CTX::*pqsA’-lux* fusion compared with the PA14 wild type (white bar) and the Δ*tatABC* mutant (black bar). The bars represent mutants where *pqsA* expression is higher (green bar) or lower (grey bar) or the same (grey bar) as the wild type. PA14 mutant 27 (pink bar) has the lowest *pqsA* expression and carries a deletion in PA14-57570 (*petA*), the Rieske subunit of cytochrome *bc_1_*. (**B**) Production of PQS, HHQ and HQNO after 16 h growth by *P. aeruginosa* PA14, the Δ*tatABC* and Δ*petA* mutants and the genetically complemented PA14 mutant (Δ*petA*::(CTX::*petA*)). ***p < 0.001, **p < 0.01, and *p < 0.05; n.s. not significant.

To confirm that the Δ*petA* mutant in common with the *P. aeruginosa tat* mutants exhibited similar defects as a consequence of perturbed PQS signalling, we compared their AQ production (**Fig. 9B**), *rhlA* expression, and eDNA release in biofilms (**S6 Fig.** and **S7 Fig.**). Comparison of PQS, HHQ and HQNO production in PA14 with the Δ*petA* mutant and the strain with a *cis* complementation of the *petA* mutation at the CTX site confirmed the loss and increase in AQ production (**Fig. 9B**). In *P. aeruginosa* PA14, the *rhlA* expression profiles for the parent and Δ*tatABC* deletion mutant in the absence or presence of exogenously supplied PQS were similar to those of strain PAO1 (compare **S6 Fig.** with **Fig. 7A**). Confocal microscopy images of biofilm formation by *P. aeruginosa* PA14 under static growth conditions shows that in common with the Δ*tatABC* mutant, *petA* mutant biofilms lack the eDNA content observed in the wild type and complemented mutant (**S7 Fig.**) demonstrating that PetA is the Tat substrate required for PQS-dependent eDNA release.

## Discussion

Extracellular DNA is one of the most abundant components of the extracellular matrix of bacterial biofilms [10–14]. However, the detailed mechanism(s) by which eDNA is released is not fully understood but likely to involve multiple pathways. *P. aeruginosa pqsL* and *pqsA* mutants release more or less eDNA respectively, consistent with the levels of PQS they produce [8]. Furthermore, the biofilms formed by *pqsA* mutants are thin and flat and contain little eDNA consistent with *pqs* signalling playing an important regulatory role in biofilm maturation [8]. To gain further insights into eDNA release, a *P. aeruginosa* transposon mutant library was screened and two groups of mutants exhibiting reduced eDNA release were identified. The first of these contained Tn insertions in genes involved in *pqs* signalling including *pqsC*, *pqsH* and *pqsR*. Mutations in *pqsC* or *pqsR* in common with *pqsA* both abrogate AQ biosynthesis while *pqsH* mutants are unable to produce 3-hydroxy-AQs such as PQS but maintain production of AQs and AQ N-oxides such as HHQ and HQNO respectively [17, 54]. These data suggest that eDNA release is likely to require PQS/HHQ rather than the effector protein PqsE. This is because although *pqsE* mutants in common with *pqsA* mutants form poor biofilms, *pqsE pqsA* double mutants could not be complemented by *pqsE* alone to restore biofilm development [26]. This contrasts with pyocyanin and lectin A for example, as production of both can be restored by PqsE in the absence of AQ biosynthesis [26].

The transposon insertions in the second group of *P. aeruginosa* eDNA release mutants obtained were located in *tatA* and *tatB*, the first two genes in the *tatABC* operon that codes for the twin-arginine translocase secretion pathway [42, 43]. The Tat secretion system was originally discovered in the thylakoid membranes of plant chloroplasts and subsequently identified in bacteria and archaea [42]. Tat exports folded proteins out of the cytoplasm and across the cytoplasmic membrane in an ATP-independent manner [42]. In bacteria such as *E. coli* and *P. aeruginosa* proteins exported by the Tat system are most frequently terminally localized in the periplasm, but some (e.g. phospholipase C) can be transported across the outer membrane by the Xcp type II secretion system, thereby becoming extracellular. Tat signal peptides harbor a conserved S/TRRxFLK consensus motif, where the twin arginine is invariant and in the vast majority of cases essential for export [45]. Using prediction algorithms for Tat substrates, 44 putative *P. aeruginosa* PA14 Tat signal peptides have been identified, and of these, 34 confirmed experimentally using a novel amidase-based Tat reporter [45]. These include phospholipases and proteins involved in pyoverdine-mediated iron-uptake, respiration, osmotic stress defence, motility, and biofilm formation [44, 45]. However, none of these are known to be involved in AQ biosynthesis, transport or *pqs* signal transduction.

In our flow-chamber grown biofilms, clear differences between the *P. aeruginosa* PAO1 wild type and *tatA* mutant biofilms were apparent. While the wild type formed characteristic mushroom-shaped structures under these conditions, the *tatA* mutant formed flat, thin biofilms. In addition, eDNA was observed primarily in the stalks of the mushroom-shaped structures in the wild-type biofilms, whereas the *tatA* mutant biofilms contained no stalks and little extracellular DNA. Moreover, the biofilms formed by the *tatA* mutant were more sensitive to tobramycin treatment than wild-type biofilms, presumably in part due to the inability of the *tatA* mutant to release DNA and form differentiated multicellular structures. Indeed, DNA binds positively charged antibiotics and can act as a ‘shield’ against aminoglycosides such as tobramycin [53]. In *P. aeruginosa* biofilms, exogenously provided DNA integrates into *P. aeruginosa* biofilms increasing their tolerance toward aminoglycosides [55]. The formation of flat, eDNA deficient, tobramycin-sensitive biofilms and the reduction in rhamnolipid, pyocyanin and MV production by the *P. aeruginosa tatA* mutant are consistent with the *pqsA* mutant biofilm phenotype [8] and imply the existence of a link between Tat export and *pqs* signalling. Since transcriptomic studies of PQS signalling in *P. aeruginosa* have not provided any evidence that the Tat export system is QS controlled [17, 26], we considered it likely that mutation of the *tat* genes resulted in perturbed PQS signalling. This observation could also account, at least in part, for the reduced virulence of *P. aeruginosa tat* mutants in a rat pulmonary infection model [44].

To explore this hypothesis, the expression of *pqsA* and the concentrations of PQS and HHQ produced were compared in *P. aeruginosa* PAO1 wild type, *tatA* and complemented *tatA* mutant strains. Similar results were also obtained for *P. aeruginosa* strain PA14. Disruption of the *tat* operon had a pronounced effect on the expression of *pqsA,* with a nearly 3–fold reduction in *pqsA* expression. Moreover, addition of the Tat inhibitor, BAY 11-7082 [46] also resulted in a comparable reduction in *pqsA* expression. LC-MS/MS quantification of PQS levels in early stationary phase after 8 h growth, showed a ∼50% reduction, thus correlating with the *pqsA’-lux* promoter fusion assays. Although PQS and HHQ both serve as PqsR ligands for driving the positive feedback loop for *pqsABCDE* expression that is central to AQ signalling [28–30], exogenous provision of neither PQS nor HHQ could fully restore *pqsA* expression in the *tatA* mutant. Since this mutant still produces AQs, albeit at a reduced level, a *pqsA* mutation was introduced into the *tatA* mutant background to examine the response to exogenous AQs in an AQ-negative genetic background (*tatA* Δ*pqsA*). The data obtained confirmed that the *P. aeruginosa tatA* Δ*pqsA* mutant responded only weakly to exogenously added PQS or HHQ, albeit to a much lesser extent than the isogenic Δ*pqsA* mutant with an intact *tat* operon. These data suggest that disruption of *tat* impairs the ability of *P. aeruginosa* to fully induce AQ-dependent QS. The consequences of *tat*-dependent perturbation of *pqs* signalling are clearly evident in the modified expression profile of the *rhlA’-lux* fusion in the *tatA* mutant with or without exogenously supplied PQS. Similar results were also obtained for *phzA1* expression in that the reduction in expression observed in a *tatA* Δ*pqsA* mutant background was not restored to wild type levels by exogenous PQS.

Since pyocyanin can be produced in the absence of PQS by ectopic expression of *pqsE* [26], it was possible that the *tat* phenotype is either a consequence of the inability of PQS/HHQ to activate the *pqsABCDE* operon via PqsR to generate sufficient PqsE protein or because the activity of PqsE depends on a functional Tat system. To investigate the impact of PqsE on pyocyanin production in the PAO1 *tatA* mutants, a plasmid borne copy of *pqsE* was introduced into the *tat* mutants. Pyocyanin production was fully restored suggesting that the *tat* mutant phenotype is not due to the inability of PqsE to function but rather a failure of the *pqs* auto-induction circuitry to produce sufficient PQS/HHQ to efficiently activate *pqsABCDE* transcription and hence *pqsE* expression.

AQs such as PQS are extremely hydrophobic and virtually insoluble in aqueous environments. Hence PQS signalling can be enhanced by increasing the solubility and delivery of PQS. Calfee *et al* [53] showed that rhamnolipids increase the solubility and bioactivity of PQS in *P. aeruginosa* with respect to *lasB* expression. Since rhamnolipid production is substantially reduced in the *P. aeruginosa tat* mutants, we added back purified rhamnolipids to the *tatA* Δ*pqsA* mutant. However, no effect on *pqsA* expression was observed indicating that the PQS signalling defect in the *tat* mutants is not simply due to the loss of rhamnolipid production and an inability to solubilize PQS.

In *Escherichia coli,* inactivation of *tat* leads to a characteristic cell envelope defect because of the mis-localization of two amidases involved in cell wall metabolism. This results in the formation of bacterial chains, leakage of periplasmic proteins and enhanced susceptibility to detergents [56]. Since PQS is a signal molecule that must be transported across the cell envelope in both directions, it was possible that an envelope defect in *P. aeruginosa tat* mutants impacts on the PQS signalling auto-induction cascade. However, Ball *et al* [57] showed that *P. aeruginosa tat* mutants do not show the same Tat-dependent envelope defects found in *E. coli.* In addition, we found no evidence for intracellular accumulation of AQs indicative of a transport defect.

To uncover the mechanistic link between Tat and PQS signalling, we sought to determine whether mutation of *tat* resulted in the inability to export a specific Tat substrate, the loss of which resulted in perturbation of PQS autoinduction and hence the expression of PQS-dependent genes and the formation of defective biofilms. By screening 34 validated Tat substrate mutants for reduced *pqsA* expression, we identified the *petA* gene as responsible for the defect in *pqs*-dependent QS. This gene codes for the Rieske protein, an iron-sulfur cluster protein sub-unit component of the cytochrome *bc_1_* complex involved in electron transfer and respiration under aerobic conditions and also essential for enabling *P. aeruginosa* to grow under anaerobic conditions in the presence of nitrite [58]. Self-poisoning of cytochrome *bc_1_* occurs via HQNO which is produced via the same biosynthetic pathway as PQS (**Fig. 1**). HQNO binds to the quinone reduction (Qi) site of the respiratory cytochrome *bc_1_* complex [59]. This results in the generation of reactive oxygen species that cause *P. aeruginosa* cell death and autolysis favouring biofilm formation and antibiotic tolerance. Rieske subunit and cytochrome *b*_1_ mutants do not undergo autolysis and are insensitive to exogenous HQNO [59]. Similarly, *P. aeruginosa pqsL* mutants that are unable to produce HQNO also fail to undergo autolysis [59]. Our data also show that in common with the *tat* mutants, the *petA* mutant produces reduced levels of HQNO (as well as PQS and HHQ). This suggests that lack of eDNA in the *tat* mutant biofilms is because the cells do not undergo limited autolysis as they lack the self-poisoning mechanism that depends on both an intact cytochrome *bc_1_* and sufficient HQNO. Thus, lack of the Rieske sub-unit export is clearly responsible for the Tat-mediated perturbation of PQS-dependent QS, the loss of virulence factor production, biofilm eDNA and the tobramycin tolerance of *P. aeruginosa* biofilms. Given the importance of PQS signalling and the Tat system to virulence and biofilm maturation in *P. aeruginosa*, our findings underline the potential of the Tat system as a drug target for novel antimicrobial agents.

## Materials and Methods

### Bacterial strains and growth conditions

The *P. aeruginosa* and *E. coli* strains used are listed in **S1 Table** and were grown in LB or ABTG [8] at 37°C unless otherwise stated. *P. aeruginosa* biofilms were cultivated at 30°C in flow-chambers irrigated with FAB medium [60] supplemented with 0.3 mM glucose or in M9 medium with succinate (for static biofilms). Selective media were supplemented with ampicillin (Ap; 100 μg ml^-1^), gentamicin (Gm; 60 μg ml^-1^), or streptomycin (Sm; 100 μg ml^-1^). PQS, HHQ and 2-heptyl-4-hydroxyquinoline N-oxide (HQNO) were synthesized and characterized in house as described before [19, 30] and dissolved in methanol before being added to growth media at the appropriate concentration.

### Mutant construction, screening and validation

The *P. aeruginosa* PAO1 Tn mutant library was constructed using the Mariner transposon vector pBT20 as previously described [61]. Transconjugants carrying Tn insertions were picked from the selective plates and inoculated into microtiter plates containing ABTG medium [62] supplemented with propidium iodide (PI) and the level of red fluorescence quantified. Mutants producing reduced levels of eDNA were selected and the sequences flanking the Tn insertion identified by arbitrary PCR essentially as described by Friedman and Kolter [63] but using the specific primers TnM1 and TnM2 (**S2 Table**). DNA Sequencing was performed by Macrogen, Seoul, Korea with primer TnMseq (**S2 Table**). An in-frame *P. aeruginosa tatABC* deletion mutant was generated by allelic exchange using the oligonucleotide primers Tat3D-UF, Tat3D-UR, Tat3D-DF and Tat3D-DR (**S2 Table**) to introduce the up- and down-stream regions of the *tatABC* locus into the suicide vector pME3087 to generate pME3087::*tatABC* (**S1 Table**). The latter was introduced into *P. aeruginosa* via conjugation with *E. coli* S17-1 λpir followed by enrichment for tetracycline-sensitive cells as described by Ye *et al* [64]. The Δ*tatABC* deletion in *P. aeruginosa* PAO1 was confirmed by PCR and sequence analysis. To generate the PA14 Δ*petA* mutant, 500 bp upstream and downstream of the gene were amplified using respectively primers S.PA4431upFor/S.PA4431upRev and S.PA4431downFor/S.PA4431downRev listed in **S2 Table**. The PCR product was cloned in pKNG101 suicide vector by one-step sequence and ligation-independent cloning (SLIC) [65] which was then sequenced. The resulting plasmid, pKNGΔ*petA*, maintained in the *E. coli* CC118λpir strain, was then mobilized in *P. aeruginosa* strains. The mutant, in which the double recombination events occurred, was confirmed by PCR analysis. A similar strategy was used to construct allelic replacement mutants for each of 34 validated Tat substrates [45]. These strains are summarized in **S3 Table** and their validation will be described in detail elsewhere. For the generation of the *petA* cis-complemented strain, PA14Δ*petA::(CTX1*::*petA)*, the *petA* genes along with a 500 bp fragment corresponding to the putative promoter region for the *petA* gene were PCR amplified using S-4431CTXFor/S-4431CTXRev and cloned by SLIC into the mini-CTX1 vector yielding pminiCTX1-*petA*. Transfer of this plasmid in *P. aeruginosa* Δ*petA* strain was carried out by triparental mating using the conjugative properties of the helper plasmid pRK2013. The recombinant clones containing the mini-CTX inserted at the *attB* locus on the *P. aeruginosa* genome were selected on tetracycline-containing PIA generating PA14Δ*petA::(attB::petA)*.

### Construction of a *tatA* complementation plasmid

The *tatA* gene was amplified by PCR using primers F*tatA* and R*tatA* (**S2 Table**), introduced into pUCP22 (**S1 Table**) and electroporated into the *P. aeruginosa tatA* and *tatA* Δ*pqsA* mutants. Transformants were selected on LB plates containing 200 μg ml^-1^ carbenicillin

### Bioluminescent reporter gene fusion assays

To investigate the impact of the *tat* mutation and Tat inhibitor Bayer 11-7082 [46] on PQS signalling, transcriptional fusions between the promoter regions of *pqsA*, *pqsR*, *rhlA*, *phzA1 and phzA2* and the *luxCDABE* operon were constructed using the miniCTX-*lux* plasmid as previously described [17]. In addition, a constitutively bioluminescent reporter using a miniCTX::*tac-luxCDABE* promoter fusion was constructed as a control for Bayer 11-7082. Bioluminescence as a function of bacterial growth was quantified in 96 well plates using a combined luminometer-spectrometer.

### Cultivation of biofilms

Biofilms were grown in flow-chambers with individual channel dimensions of 1 x 4 x 40 mm as described previously [66]. One hour after inoculation, with bacteria, the growth medium flow (0.2 mm/s corresponding to laminar flow with a Reynolds number of 0.02) was started. When required, eDNA in biofilms was stained with 1 µM ethidium bromide prior to microscopy. Tobramycin (10 µM) was added to the biofilm medium after 4 days of cultivation. After 24 h of tobramycin treatment, propidium iodide (10 µM) was added to the flow cells to visualize the dead cells via confocal laser scanning microscopy. *P. aeruginosa* PA14 and *petA* mutant biofilms were grown under static conditions over 48 h at 37 °C on glass slides (Ibidi) incorporating 300 µl chambers. After 48 h incubation, spent medium was removed and the biofilm eDNA stained with YoYo-1 (40 µM).

### Microscopy and image processing of flow cell biofilms

All images of flow-chamber-grown and static biofilms were captured with a confocal laser scanning microscope (CLSM) equipped with detectors and filter sets for monitoring green fluorescent protein, Syto9, propidium iodide, and ethidium bromide. Images were obtained using a 63x/1.4 objective or a 40x/1.3 objective. Simulated 3-D images and sections were generated using the IMARIS software package (Bitplane AG, Zürich, Switzerland).

### AQ, pyocyanin, rhamnolipid and MV analysis

The AQs (PQS, HHQ and HQNO) were quantified by LC-MS/MS after extracting cell free supernatants or whole bacterial cells in triplicate with acidified ethyl acetate or methanol respectively as described by Ortori *et al* [19]. Pyocyanin was extracted with chloroform and quantified spectrophotometrically [26]. Rhamnolipids were quantified indirectly using the orcinol method [35]. For PQS solubilization experiments, rhamnolipids were purified as described by Muller *et al* [67]. MVs were harvested by ultracentrifugation, the pellets resuspended in 10mM HEPES buffer and the lipid content quantified using FM4-64 essentially as described previously [68]. MV production was normalized by dividing the lipid fluorescence units by CFU values determined by dilution plating. Assays were performed in triplicate at least twice.

### Statistical Analysis

Significance for differences between wild type and isogenic mutants was determined by two-tailed *t*-tests where ****p < 0.001, ***p < 0.001, **p < 0.01, and *p < 0.05 and n.s., not significant.

## Supporting information

Manuscript

## Supporting Information

**S1 Fig.** The Tat inhibitor Bayer 11-7082 has no effect on light output at 20 or 40 μM from *P. aeruginosa* CTX:*ptac’-luxCDABE* chromosomal reporter fusion. The co-solvent DMSO, had no effect at 0.8% on the *lux* reporter fusion either. Data are presented as maximal light output as a function of growth (RLU/OD_600_). Experiments were repeated in triplicate at least twice.

**S2 Fig**. Deletion of the *P. aeruginosa tatABC* genes does not influence *pqsR* expression. The data show that there are no differences in the expression of a chromosomal CTX::*pqsR’-luxCDABE* fusion in the *P. aeruginosa* wild type compared with the Δ*tatABC* mutant Data are presented as maximal light output as a function of growth (RLU/OD_600_). Experiments were repeated in triplicate at least twice.

**S3 Fig.** AQ biosynthesis is not restored in a *tatA* Δ*pqsA* double mutant by plasmid-borne *pqsABCD* in the absence of autoinduction. Semi-quantitative analysis by LC-MS/MS of PQS, HHQ and HQNO production by *P. aeruginosa pqsA* and *tatA* Δ*pqsA* mutants respectively without or with the *pqsABCD* biosynthetic genes provided *in trans* via pBBRMCS::*pqsABCD.* Experiments were repeated in triplicate at least twice.

**S4 Fig.** HHQ and PQS do not accumulate intracellularly a *tatA* Δ*pqsA* double mutant harboring the plasmid-borne *pqsABCD* genes in the absence of autoinduction. Semi-quantitative analysis by LC-MS/MS of PQS (**A**) and HHQ (**B**) extracted from whole cells of *P. aeruginosa* wild type and the *tatA* Δ*pqsA* mutant without or with the *pqsABCD* biosynthetic genes provided via pBBRMCS::*pqsABCD.* Cells were harvested at 8 h and 16 h respectively. Experiments were repeated in triplicate

**S5 Fig.** Exogenous rhamnolipids do not enhance PQS-dependent expression of *pqsA* in a *tatA* Δ*pqsA* mutant. PQS (40 μM) was added with or without purified rhamnolipids (50 μg/ml) to a *pqsA* mutant or a *tatA* Δ*pqsA* mutant carrying chromosomal *pqsA’-lux* fusions. Maximal light output as a function of growth (RLU/OD_600_) is presented. Experiments were repeated in triplicate at least twice.

**S6 Fig** Rhamnolipid biosynthesis gene *rhlA* shows altered expression profiles in *P. aeruginosa* PA14 Δ*tatABC* and Δ*petA* mutants compared with wild type and fail to respond to exogenous PQS. Bioluminescence from a chromosomal *rhlA’-lux* fusion as a function of growth (RLU/OD) over time when introduced into (**A**) the PA14 wild type, (**B**) Δ*tatABC* and (**C**) Δ*petA* mutants in the absence or presence of exogenous PQS (20 μM).

**S7 Fig** Deletion of *petA* in *P. aeruginosa* PA14 results in a reduction in the eDNA content of biofilms. Biofilms of wild type PA14, the Δ*tatABC* and Δ*petA* mutants and the genetically complemented PA14 mutant (Δ*petA*::(CTX::*petA*)) were grown cultured statically and stained for eDNA with YOYO-1. (**A**) CLSM images and (**B**) eDNA quantification. Experiments were repeated in triplicate at least twice. ***p < 0.001, **p < 0.01.

**S1 Table** Strains and plasmids used in this study

**S2 Table** Oligonucleotide primers used in this study

**S3 Table** *P. aeruginosa* PA14 Tat substrate mutants used in this study

## Acknowledgements

We thank Alex Truman for synthesis of AQ standards. This work was funded via grants to PW and MC from the Biotechnology and Biological Sciences Research Council, UK (BB/F014392/1; www.bbsrc.ac.uk), the Medical Research Council U.K.(MR/N501852/1; www.mrc.ac.uk), the Wellcome Trust (103884/Z/14/Z; www.welcome.ac.uk), the European Union FP7 collaborative action grant (NABATIVI, Ref. 223670; http://cordis.europa.eu/project/rcn/90762_en.html). MC, PW and KH are partly funded by the National Biofilms Innovation Centre (NBIC) which is an Innovation and Knowledge Centre funded by the BBSRC, InnovateUK and Hartree Centre (Award Number BB/R012415/1). FA was funded via a Wellcome Trust Doctoral Training Programme (108876/Z/15/Z). BI was funded by a grant from Vaincre la Mucoviscidose and the Grégory Lemarchal associations (RF20140501138) and recurrent funding from the CNRS and Aix-Marseille Université. LY, MG and TTN were funded by grants from the Danish Strategic Research Council, the Danish Council for Independent Research, the Novo Nordisk Foundation and the Lundbeck Foundation. MRG was supported by a PhD studentship and ATER position from Aix-Marseille Université.

## Notes

### Competing Interest Statement

The authors have declared no competing interest.

### Summary of Updates

Supplementary Information Added

## References

1. Moradali MF, Ghods S, Rehm BH. *Pseudomonas aeruginosa* Lifestyle: A paradigm for adaptation, survival, and persistence. Front Cell Infect Microbiol. 2017; 7:39–42. doi: 10.3389/fcimb.2017.00039. PMID: 28261568

2. Costerton JW, Stewart PS, Greenberg EP. Bacterial biofilms: a common cause of persistent infections. Science. 1999; 284:1318–22. doi: 10.1126/science.284.5418.1318. PMID: 10334980.

3. Klausen M, Aaes-Jørgensen A, Molin S, Tolker-Nielsen T. Involvement of bacterial migration in the development of complex multicellular structures in *Pseudomonas aeruginosa* biofilms. Mol Microbiol. 2003; 50:61–8. doi: 10.1046/j.1365-2958.2003.03677.x. PMID: 14507363.

4. Klausen M, Heydorn A, Ragas P, Lambertsen L, Aaes-Jørgensen A, Molin S, Tolker-Nielsen T. Biofilm formation by *Pseudomonas aeruginosa* wild type, flagella and type IV pili mutants. Mol Microbiol. 2003; 48:1511–24. doi: 10.1046/j.1365-2958.2003.03525.x. PMID: 12791135.

5. Pamp SJ, Tolker-Nielsen T. Multiple roles of biosurfactants in structural biofilm development by *Pseudomonas aeruginosa*. J Bacteriol. 2007; 189:2531–39. doi: 10.1128/JB.01515-06. PMID: 17220224

6. Harmsen M, Yang L, Pamp SJ, Tolker-Nielsen T. An update on *Pseudomonas aeruginosa* biofilm formation, tolerance, and dispersal. FEMS Immunol Med Microbiol. 2010; 59:253–68. doi: 10.1111/j.1574-695X.2010.00690.x. PMID: 20497222.

7. Yang L, Hu Y, Liu Y, Zhang J, Ulstrup J, Molin S. Distinct roles of extracellular polymeric substances in *Pseudomonas aeruginosa* biofilm development. Environ Microbiol. 2011; 13:1705–17. doi: 10.1111/j.1462-2920.2011.02503.x. PMID: 21605307.

8. Allesen-Holm M, Barken KB, Yang L, Klausen M, Webb JS, Kjelleberg S, Molin S, Givskov M, Tolker-Nielsen T. A characterization of DNA release in *Pseudomonas aeruginosa* cultures and biofilms. Mol Microbiol. 2006; 59:1114–28. doi: 10.1111/j.1365-2958.2005.05008.x. PMID: 16430688.

9. Tashiro Y, Uchiyama H, Nomura N. Multifunctional membrane vesicles in *Pseudomonas aeruginosa*. Environ Microbiol. 2012; 14:1349–62. doi: 10.1111/j.1462-2920.2011.02632.x. PMID: 22103313.

10. Tolker-Nielsen T. Biofilm Development. Microbiol Spectr. 2015 3:MB-0001-2014. doi: 10.1128/microbiolspec.MB-0001-2014. PMID: 26104692.

11. Okshevsky M, Meyer RL. The role of extracellular DNA in the establishment, maintenance and perpetuation of bacterial biofilms. Crit Rev Microbiol. 2015;41:341–52. doi: 10.3109/1040841X.2013.841639. PMID: 24303798.

12. Das T, Sehar S, Manefield M. The roles of extracellular DNA in the structural integrity of extracellular polymeric substance and bacterial biofilm development. Environ Microbiol Rep. 2013 5:778–86. doi: 10.1111/1758-2229.12085. PMID: 24249286.

13. Wilton M, Charron-Mazenod L, Moore R, Lewenza S. Extracellular DNA acidifies biofilms and induces aminoglycoside resistance in *Pseudomonas aeruginosa*. Antimicrob Agents Chemother. 2015; 60:544–53. doi: 10.1128/AAC.01650-15. PMID: 26552982

14. Turnbull L, Toyofuku M, Hynen AL, Kurosawa M, Pessi G, Petty NK, Osvath SR, Cárcamo-Oyarce G, Gloag ES, Shimoni R, Omasits U, Ito S, Yap X, Monahan LG, Cavaliere R, Ahrens CH, Charles IG, Nomura N, Eberl L, Whitchurch CB. Explosive cell lysis as a mechanism for the biogenesis of bacterial membrane vesicles and biofilms. Nat Commun. 2016; 7:11220. doi: 10.1038/ncomms11220. PMID: 27075392.

15. Das T, Kutty SK, Tavallaie R, Ibugo AI, Panchompoo J, Sehar S, Aldous L, Yeung AW, Thomas SR, Kumar N, Gooding JJ, Manefield M. Phenazine virulence factor binding to extracellular DNA is important for *Pseudomonas aeruginosa* biofilm formation. Sci Rep. 2015; 5:8398. doi: 10.1038/srep08398. PMID: 25669133.

16. Williams P, Cámara M. Quorum sensing and environmental adaptation in *Pseudomonas aeruginosa*: a tale of regulatory networks and multifunctional signal molecules. Curr Opin Microbiol. 2009; 12:182–91. doi: 10.1016/j.mib.2009.01.005. PMID: 19249239.

17. Rampioni G, Falcone M, Heeb S, Frangipani E, Fletcher MP, Dubern JF, Visca P, Leoni L, Cámara M, Williams P. Unravelling the genome-wide contributions of specific 2-alkyl-4-quinolones and PqsE to quorum sensing in *Pseudomonas aeruginosa*. PLoS Pathog. 2016; 12:e1006029. doi: 10.1371/journal.ppat.1006029. PMID: 27851827.

18. Lépine F, Milot S, Déziel E, He J, Rahme LG. Electrospray/mass spectrometric identification and analysis of 4-hydroxy-2-alkylquinolines (HAQs) produced by *Pseudomonas aeruginosa*. J Am Soc Mass Spectrom. 2004; 15:862–9. doi: 10.1016/j.jasms.2004.02.012. PMID: 15144975.

19. Ortori CA, Dubern JF, Chhabra SR, Cámara M, Hardie K, Williams P, Barrett DA. Simultaneous quantitative profiling of *N*-acyl-L-homoserine lactone and 2-alkyl-4(1*H*)-quinolone families of quorum-sensing signaling molecules using LC-MS/MS. Anal Bioanal Chem. 2011; 399:839–50. doi: 10.1007/s00216-010-4341-0. PMID: 21046079.

20. Coleman JP, Hudson LL, McKnight SL, Farrow JM 3rd, Calfee MW, Lindsey CA, Pesci EC. *Pseudomonas aeruginosa* PqsA is an anthranilate-coenzyme A ligase. J Bacteriol. 2008; 190:1247–55. doi: 10.1128/JB.01140-07. PMID: 18083812

21. Zhang YM, Frank MW, Zhu K, Mayasundari A, Rock CO. PqsD is responsible for the synthesis of 2,4-dihydroxyquinoline, an extracellular metabolite produced by *Pseudomonas aeruginosa*. J Biol Chem. 2008; 283:28788–94. doi: 10.1074/jbc.M804555200. PMID: 18728009.

22. Drees SL, Fetzner S. PqsE of *Pseudomonas aeruginosa* acts as pathway-specific thioesterase in the biosynthesis of alkylquinolone signaling molecules. Chem Biol. 2015; 22:611–8. doi: 10.1016/j.chembiol.2015.04.012. PMID: 25960261.

23. Dulcey CE, Dekimpe V, Fauvelle DA, Milot S, Groleau MC, Doucet N, Rahme LG, Lépine F, Déziel E. The end of an old hypothesis: the pseudomonas signaling molecules 4-hydroxy-2-alkylquinolines derive from fatty acids, not 3-keto fatty acids. Chem Biol. 2013; 20:1481–91. doi: 10.1016/j.chembiol.2013.09.021. PMID: 24239007

24. Drees SL, Li C, Prasetya F, Saleem M, Dreveny I, Williams P, Hennecke U, Emsley J, Fetzner S. PqsBC, a condensing enzyme in the biosynthesis of the *Pseudomonas aeruginosa* quinolone signal: crystal structure, inhibition, and reaction mechanism. J Biol Chem. 2016; 291:6610–24. doi: 10.1074/jbc.M115.708453. PMID: 26811339;

25. Schertzer JW, Brown SA, Whiteley M. Oxygen levels rapidly modulate *Pseudomonas aeruginosa* social behaviours via substrate limitation of PqsH. Mol Microbiol. 2010; 77:1527–38. doi: 10.1111/j.1365-2958.2010.07303.x. PubMed PMID: 20662781.

26. Rampioni G, Pustelny C, Fletcher MP, Wright VJ, Bruce M, Rumbaugh KP, Heeb S, Cámara M, Williams P. Transcriptomic analysis reveals a global alkyl-quinolone-independent regulatory role for PqsE in facilitating the environmental adaptation of *Pseudomonas aeruginosa* to plant and animal hosts. Environ Microbiol. 2010; 12:1659–73. doi: 10.1111/j.1462-2920.2010.02214.x. PMID: 20406282.

27. Groleau, MC, de Oliveira Pereira, T, Dekimpe V, Déziel E. PqsE Is Essential for RhlR-Dependent Quorum Sensing Regulation in *Pseudomonas aeruginosa* mSystems. 2020; 5:e00194–20; doi: 10.1128/mSystems.00194-20. PMID: 32457239

28. Wade DS, Calfee MW, Rocha ER, Ling EA, Engstrom E, Coleman JP, Pesci EC. Regulation of pseudomonas quinolone signal synthesis in *Pseudomonas aeruginosa*. J Bacteriol. 2005;187:4372–80. doi: 10.1128/JB.187.13.4372-4380.2005. PMID: 15968046.

29. Xiao G, Déziel E, He J, Lépine F, Lesic B, Castonguay MH, Milot S, Tampakaki AP, Stachel SE, Rahme LG. MvfR, a key *Pseudomonas aeruginosa* pathogenicity LTTR-class regulatory protein, has dual ligands. Mol Microbiol. 2006; 62:1689–99. doi: 10.1111/j.1365-2958.2006.05462.x. PMID: 17083468.

30. Ilangovan A, Fletcher M, Rampioni G, Pustelny C, Rumbaugh K, Heeb S, Cámara M, Truman A, Chhabra SR, Emsley J, Williams P. Structural basis for native agonist and synthetic inhibitor recognition by the *Pseudomonas aeruginosa* quorum sensing regulator PqsR (MvfR). PLoS Pathog. 2013; 9:e1003508. doi: 10.1371/journal.ppat.1003508. PMID: 23935486.

31. Diggle SP, Matthijs S, Wright VJ, Fletcher MP, Chhabra SR, Lamont IL, Kong X, Hider RC, Cornelis P, Cámara M, Williams P. The *Pseudomonas aeruginosa* 4-quinolone signal molecules HHQ and PQS play multifunctional roles in quorum sensing and iron entrapment. Chem Biol. 2007;14:87–96. doi: 10.1016/j.chembiol.2006.11.01. PMID: 17254955.

32. Häussler S, Becker T. The pseudomonas quinolone signal (PQS) balances life and death in *Pseudomonas aeruginosa* populations. PLoS Pathog. 2008; 4:e1000166. doi: 10.1371/journal.ppat.1000166. PMID: 18818733;

33. Mashburn-Warren L, Howe J, Garidel P, Richter W, Steiniger F, Roessle M, Brandenburg K, Whiteley M. Interaction of quorum signals with outer membrane lipids: insights into prokaryotic membrane vesicle formation. Mol Microbiol. 2008 69:491–502. doi: 10.1111/j.1365-2958.2008.06302.x. PMID: 18630345.

34. Mashburn LM, Whiteley M. Membrane vesicles traffic signals and facilitate group activities in a prokaryote. Nature. 2005; 437:422–5. doi: 10.1038/nature0392. PMID: 16163359

35. Pustelny C, Albers A, Büldt-Karentzopoulos K, Parschat K, Chhabra SR, Cámara M, Williams P, Fetzner S. Dioxygenase-mediated quenching of quinolone-dependent quorum sensing in *Pseudomonas aeruginosa*. Chem Biol. 2009; 16:1259–67. doi: 10.1016/j.chembiol.2009.11.013. PMID: 20064436.

36. D’Argenio DA, Calfee MW, Rainey PB, Pesci EC. Autolysis and autoaggregation in *Pseudomonas aeruginosa* colony morphology mutants. J Bacteriol. 2002; 184:6481–9. doi: 10.1128/jb.184.23.6481-6489.2002. PMID: 12426335.

37. Webb JS, Thompson LS, James S, Charlton T, Tolker-Nielsen T, Koch B, Givskov M, Kjelleberg S. Cell death in *Pseudomonas aeruginosa* biofilm development. J Bacteriol. 2003; 185:4585–92. doi: 10.1128/jb.185.15.4585-4592.200. PMID: 12867469

38. Spoering AL, Gilmore MS. Quorum sensing and DNA release in bacterial biofilms. Curr Opin Microbiol. 2006; 9:133–7. doi: 10.1016/j.mib.2006.02.004. PMID: 16529982.

39. Yang L, Barken KB, Skindersoe ME, Christensen AB, Givskov M, Tolker-Nielsen T. Effects of iron on DNA release and biofilm development by *Pseudomonas aeruginosa*. Microbiology. 2007;153:1318–28. doi: 10.1099/mic.0.2006/004911-0. PMID: 17464046.

40. Yang L, Nilsson M, Gjermansen M, Givskov M, Tolker-Nielsen T. Pyoverdine and PQS mediated subpopulation interactions involved in *Pseudomonas aeruginosa* biofilm formation. Mol Microbiol. 2009;74:1380–92. doi: 10.1111/j.1365-2958.2009.06934.x. PMID: 19889094.

41. Suzuki T, Fujikura K, Higashiyama T, Takata K. DNA staining for fluorescence and laser confocal microscopy. J Histochem Cytochem. 1997; 45:49–53. doi: 10.1177/002215549704500107. PMID: 9010468.

42. Patel R, Smith SM, Robinson C. Protein transport by the bacterial Tat pathway. Biochim Biophys Acta. 2014; 1843:1620–8. doi: 10.1016/j.bbamcr.2014.02.013. PMID: 24583120.

43. Costa TR, Felisberto-Rodrigues C, Meir A, Prevost MS, Redzej A, Trokter M, Waksman G. Secretion systems in Gram-negative bacteria: structural and mechanistic insights. Nat Rev Microbiol. 2015; 13:343–59. doi: 10.1038/nrmicro3456. PMID: 25978706.

44. Ochsner UA, Snyder A, Vasil AI, Vasil ML. Effects of the twin-arginine translocase on secretion of virulence factors, stress response, and pathogenesis. Proc Natl Acad Sci U S A. 2002; 99:8312–7. doi: 10.1073/pnas.082238299. PMID: 12034867.

45. Gimenez MR, Chandra G, Van Overvelt P, Voulhoux R, Ize, B, Bleves, S. Genome wide identification and experimental validation of *Pseudomonas aeruginosa* Tat substrates. Sci Rep 2018; 11950. doi: 10.1038/s41598-018-30393-x. PMID: 30093651

46. Vasil ML, Tomaras AP, Pritchard AE. Identification and evaluation of twin-arginine translocase inhibitors. Antimicrob Agents Chemother. 2012; 56:6223–34. doi: 10.1128/AAC.01575-12. PMID: 23006747

47. Niewerth H, Bergander K, Chhabra SR, Williams P, Fetzner S. Synthesis and biotransformation of 2-alkyl-4(1H)-quinolones by recombinant *Pseudomonas putida* KT2440. Appl Microbiol Biotechnol. 2011; 91:1399–408. doi: 10.1007/s00253-011-3378-0. PMID: 21670979.

48. Williams P, Winzer K, Chan WC, Cámara M. Look who’s talking: communication and quorum sensing in the bacterial world. Philos Trans R Soc Lond B Biol Sci. 2007; 362:1119–34. doi: 10.1098/rstb.2007.2039. PMID: 17360280;

49. Recinos DA, Sekedat MD, Hernandez A, Cohen TS, Sakhtah H, Prince AS, Price-Whelan A, Dietrich LE. Redundant phenazine operons in *Pseudomonas aeruginosa* exhibit environment-dependent expression and differential roles in pathogenicity. Proc Natl Acad Sci U S A. 2012; 109:19420–5. doi: 10.1073/pnas.1213901109. PMID: 23129634

50. Farrow JM 3rd, Sund ZM, Ellison ML, Wade DS, Coleman JP, Pesci EC. PqsE functions independently of PqsR-Pseudomonas quinolone signal and enhances the *rhl* quorum-sensing system. J Bacteriol. 2008; 190:7043–51. doi: 10.1128/JB.00753-08. PMID: 18776012

51. Van Gennip M, Christensen LD, Alhede M, Phipps R, Jensen PØ, Christophersen L, Pamp SJ, Moser C, Mikkelsen PJ, Koh AY, Tolker-Nielsen T, Pier GB, Høiby N, Givskov M, Bjarnsholt T. Inactivation of the *rhlA* gene in *Pseudomonas aeruginosa* prevents rhamnolipid production, disabling the protection against polymorphonuclear leukocytes. APMIS. 2009 117:537–46. doi: 10.1111/j.1600-0463.2009.02466.x. PMID: 19594494

52. Alhede M, Bjarnsholt T, Jensen PØ, Phipps RK, Moser C, Christophersen L, Christensen LD, van Gennip M, Parsek M, Høiby N, Rasmussen TB, Givskov M. *Pseudomonas aeruginosa* recognizes and responds aggressively to the presence of polymorphonuclear leukocytes. Microbiology (Reading). 2009 155:3500–3508. doi: 10.1099/mic.0.031443-0. PMID: 19643762.

53. Calfee MW, Shelton JG, McCubrey JA, Pesci EC. Solubility and bioactivity of the Pseudomonas quinolone signal are increased by a *Pseudomonas aeruginosa*-produced surfactant. Infect Immun. 2005; 73:878–82. doi: 10.1128/IAI.73.2.878-882.2005. PMID: 15664929.

54. Déziel E, Lépine F, Milot S, He J, Mindrinos MN, Tompkins RG, Rahme LG. Analysis of *Pseudomonas aeruginosa* 4-hydroxy-2-alkylquinolines (HAQs) reveals a role for 4-hydroxy-2-heptylquinoline in cell-to-cell communication. Proc Natl Acad Sci U S A. 2004; 101:1339–44. doi: 10.1073/pnas.0307694100. PMID: 14739337

55. Chiang WC, Nilsson M, Jensen PØ, Høiby N, Nielsen TE, Givskov M, Tolker-Nielsen T. Extracellular DNA shields against aminoglycosides in *Pseudomonas aeruginosa* biofilms. Antimicrob Agents Chemother. 2013; 57:2352–61. doi: 10.1128/AAC.00001-13. PMID: 23478967

56. Ize B, Stanley NR, Buchanan G, Palmer T. Role of the *Escherichia coli* Tat pathway in outer membrane integrity. Mol Microbiol. 2003; 48:1183–93. doi: 10.1046/j.1365-2958.2003.03504.x.PMID: 12787348.

57. Ball G, Antelmann H, Imbert PR, Gimenez MR, Voulhoux R, Ize B. Contribution of the twin arginine translocation system to the exoproteome of *Pseudomonas aeruginosa*. Sci Rep. 2016; 6:27675. doi: 10.1038/srep27675. PMID: 27279369

58. Hasegawa N, Arai H, Igarashi Y. Need for Cytochrome bc 1 Complex for dissimilatory nitrite reduction of *Pseudomonas aeruginosa*. Biosci Biotechnol Biochem. 2003 67:121–126. doi: 10.1271/bbb.67.12. PMID: 12619683.

59. Hazan R, Que YA, Maura D, Strobel B, Majcherczyk PA, Hopper LR, Wilbur DJ, Hreha TN, Barquera B, Rahme LG. Auto poisoning of the respiratory chain by a quorum-sensing-regulated molecule favors biofilm formation and antibiotic tolerance. Curr Biol. 2016; 26:195–206. doi: 10.1016/j.cub.2015.11.056. PMID: 26776731

60. Heydorn A, Nielsen AT, Hentzer M, Sternberg C, Givskov M, Ersbøll BK, Molin S. Quantification of biofilm structures by the novel computer program COMSTAT. Microbiology. 2000;146:2395–407 doi: 10.1099/00221287-146-10-2395. PMID: 11021916.

61. Kulasekara HD, Ventre I, Kulasekara BR, Lazdunski A, Filloux A, Lory S. A novel two-component system controls the expression of *Pseudomonas aeruginosa* fimbrial cup genes. Mol Microbiol. 2005 Jan;55(2):368–80. doi: 10.1111/j.1365-2958.2004.04402.x. PMID: 15659157.

62. Chua SL, Tan SY, Rybtke MT, Chen Y, Rice SA, Kjelleberg S, Tolker-Nielsen T, Yang L, Givskov M. Bis-(3’-5’)-cyclic dimeric GMP regulates antimicrobial peptide resistance in *Pseudomonas aeruginosa*. Antimicrob Agents Chemother. 2013; 57:2066–75. doi: 10.1128/AAC.02499-12. PMID: 23403434.

63. Friedman L, Kolter R. Genes involved in matrix formation in *Pseudomonas aeruginosa* PA14 biofilms. Mol Microbiol. 2004; 51:675–90. doi: 10.1046/j.1365-2958.2003.03877.x. PMID: 14731271.

64. Ye RWl, Haas D, Ka JO, Krishnapillai V, Zimmermann A, Baird C, Tiedje JM. Anaerobic activation of the entire denitrification pathway in *Pseudomonas aeruginosa* requires Anr, an analog of Fnr. J Bacteriol. 1995; 177:3606–9. doi: 10.1128/jb.177.12.3606-3609.1995. PMID: 7768875

65. Jeong JY, Yim HS, Ryu JY, Lee HS, Lee JH, Seen DS, Kang SG. One-step sequence- and ligation-independent cloning as a rapid and versatile cloning method for functional genomics studies. Appl Environ Microbiol. 2012; 78:5440–3. doi: 10.1128/AEM.00844-12. PMID: 22610439.

66. Crusz SA, Popat R, Rybtke MT, Cámara M, Givskov M, Tolker-Nielsen T, Diggle SP, Williams P. Bursting the bubble on bacterial biofilms: a flow cell methodology. Biofouling. 2012; 28:835–42. doi: 10.1080/08927014.2012.716044. PMID: 22877233

67. Müller MM, Hörmann B, Syldatk C, Hausmann R. *Pseudomonas aeruginosa* PAO1 as a model for rhamnolipid production in bioreactor systems. Appl Microbiol Biotechnol. 2010; 87:167–74. doi: 10.1007/s00253-010-2513-7. PMID: 20217074.

68. McBroom AJ, Johnson AP, Vemulapalli S, Kuehn MJ. Outer membrane vesicle production by *Escherichia coli* is independent of membrane instability. J Bacteriol. 2006; 188:5385–92. doi: 10.1128/JB.00498-06. PMID: 16855227.

