## Supplementary material for "Disruption of the *Pseudomonas aeruginosa* Tat system perturbs PQS-dependent quorum sensing and biofilm maturation through loss of the Rieske cytochrome *bc_1_* sub-unit": Manuscript

### Supporting Information

**S1 Fig.** The Tat inhibitor Bayer 11-7082 has no effect on light output at 20 or 40 µM from *P. aeruginosa* CTX:*ptac'-luxCDABE* chromosomal reporter fusion.

**S2 Fig**. Deletion of the *P. aeruginosa tatABC* genes does not influence *pqsR* expression.

**S3 Fig.** AQ biosynthesis is not restored in a *tatA* Δ*pqsA* double mutant by plasmid-borne *pqsABCD* in the absence of autoinduction.

**S4 Fig.** HHQ and PQS do not accumulate intracellularly a *tatA* Δ*pqsA* double mutant harboring the plasmid-borne *pqsABCD* genes in the absence of autoinduction.

**S5 Fig.** Exogenous rhamnolipids do not enhance PQS-dependent expression of *pqsA* in a *tatA* Δ*pqsA* mutant.

**S6 Fig** Rhamnolipid biosynthesis gene *rhlA* shows altered expression profiles in *P. aeruginosa* PA14 Δ*tatABC* and Δ*petA* mutants compared with wild type and fail to respond to exogenous PQS.

**S7 Fig** Deletion of *petA* in *P. aeruginosa* PA14 results in a reduction in the eDNA content of biofilms.

**S1 Table** Strains and plasmids used in this study

**S2 Table** Oligonucleotide primers used in this study

**S3 Table** *P. aeruginosa* PA14 Tat substrate mutants used in this study

**Table S1.** Strains and plasmids used in this study

| Strains/Plasmids | Relevant Characteristics | Source/Reference |
|---|---|---|
| ***E. coli*** | | |
| S17-1λ*pir* | Conjugative strain for suicide plasmids | [1] |
| ***P. aeruginosa*** | | |
| PAO1 | Wild-type *P. aeruginosa* strain | Copenhagen collection |
| PAO1 CTX::*pqsA'-lux* | PAO1 with a chromosomal miniCTX::*pqsA'-luxCDABE* fusion | [2] |
| PAO1 CTX::*rhlA'-lux* | PAO1 with a chromosomal miniCTX::*rhlA'-luxCDABE* fusion | This study |
| PAO1 CTX::*phzA1'-lux* | PAO1 with a chromosomal miniCTX::*phzA1'-luxCDABE* fusion integrated into the chromosomal CTX attachment site | [3] |
| PAO1 CTX::*phzA2'-lux* | PAO1 with a chromosomal miniCTX::*phzA2'-luxCDABE* fusion | [3] |
| PAO1 CTX::*pqsR'-lux* | PAO1 with a chromosomal miniCTX::*pqsR'-luxCDABE* fusion | This study |
| PAO1 CTX::*tac-lux* | PAO1 with a chromosomal miniCTX::*tac'-luxCDABE* fusion | This study |
| PAO1 Δ*pqsA* | *pqsA* in-frame deletion mutant; AQ-negative | [2] |
| PAO1 Δ*pqsA* CTX::*pqsA'-lux* | PAO1 Δ*pqsA* with a chromosomal miniCTX::*pqsA'-luxCDABE* fusion | [2] |
| PAO1 Δ*pqsA* CTX::*rhlA'-lux* | PAO1 Δ*pqsA* with a chromosomal miniCTX::*rhlA'-luxCDABE* fusion | This study |
| PAO1 *tatA* | *tatA* Himar 1 *mariner* transposon insertion mutant | This study |
| PAO1 Δ*tatABC* | *tatABC* in-frame deletion mutant | This study |
| PAO1 *tatA* Δ*pqsA* | PAO1 *tatA* with a *pqsA* in-frame deletion | This study |
| PAO1 *tatA* CTX::*pqsA'-lux* | PAO1 *tatA* with a chromosomal miniCTX::*pqsA'-luxCDABE* fusion | This study |
| PAO1 *tatA* CTX::*rhlA'-lux* | PAO1 *tatA* with a chromosomal miniCTX::*rhlA'-luxCDABE* fusion | This study |
| PAO1 *tatA* CTX::*phzA1'-lux* | PAO1 *tatA* with a chromosomal miniCTX::*phzA1'-luxCDABE* fusion | This study |
| PAO1 *tatA* CTX::*phzA2'-lux* | PAO1 *tatA* with a chromosomal miniCTX::*phzA2'-luxCDABE* fusion | This study |
| PAO1 *tatA* Δ*pqsA* CTX::*pqsA'-lux* | PAO1 *tatA* Δ*pqsA* with a chromosomal miniCTX::*pqsA'-luxCDABE* fusion | This study |
| PA14 | Wild-type *P. aeruginosa* strain | This study |
| PA14 Δ*petA* | *petA* in-frame deletion mutant | This study |
| PA14 Δ*petA*::(CTX1::*petA*) | *petA* with a chromosomal miniCTX1*::petA* insertion, $Tc^R$ | This study |
| **Plasmids** | | |
| pBT20 | *Himar I mariner* mini-transposon delivery vector | [4] |
| pME3087 | ColE1 suicide vector for allelic replacements, $Tc^R$ | [5] |
| pBBR1MCS-5 | Broad host range vector, $Gm^R$ | [6] |
| pBBR1MCS-5::*pqsABCD* | pBBR1MCS-5 carrying the *pqsABCD* operon | [7] |
| pME6032::*pqsR6H* | pME6032 carrying a functional, hexahistidine C-terminally tagged *pqsR gene* | [8] |
| pUCP22 | *E.coli -Pseudomonas* shuttle vector | [9] |
| pTatA | pUCP22 carrying *tatA*; $Ap^R$ | This study |
| pUCP*pqsE* | pUCP18 containing *pqsE*; $Ap^R$ | [10] |
| pmini-CTX-*lux* | Promoter probe vector containing *luxCDABE*, $Tc^R$ | [11] |
| pminiCTX::*pqsA'-lux* | *pqsA* promoter region fused to *luxCDABE* in pminiCTX-*lux* | [2] |
| pminiCTX::*rhlA'-lux* | *rhlA* promoter region fused to *luxCDABE* in pminiCTX-*lux* | This study |
| pminiCTX::*phzA1'-lux* | *phzA1* promoter region fused to *luxCDABE* in pminiCTX-*lux* | [3] |
| pminiCTX::*phzA2'-lux* | *phzA2* promoter region fused to *luxCDABE* in pminiCTX-*lux* | [3] |
| pminiCTX::*pqsR'-lux* | *pqsR* promoter region fused to *luxCDABE* in pminiCTX-*lux* | This study |
| pminiCTX::*tac-lux* | *tac* promoter fused to *luxCDABE* in pminiCTX-*lux* | This study |
| pRK2013 | ColE1 origin, $Tra^+$, $mob^+$, $Km^R$ | [12] |
| pKNG101 | $Sm^R$, *oriR6K, oriTRK2, mobRK2, sacBR*$^+$ | [13] |
| mini-CTX1 | Contains *attP* site for integration at the *attB* site of *P. aeruginosa* chromosome; $Tc^R$ | [14] |
| pKNGΔ*petA* | Suicide vector for *petA* deletion, $Sm^R$ | This study |
| pminiCTX1-*petA* | *petA* under the control of its own promoter in mini-CTX1 | This study |

### References for Table S1

1. Simon R, Priefer U, Puhler A. A broad host range mobilization system for *in vivo* genetic engineering: transposon mutagenesis in Gram-negative bacteria. Nature Biotechnology. 1983; 1:784-91.

2. Diggle SP, Matthijs S, Wright VJ, Fletcher MP, Chhabra SR, Lamont IL, Kong X, Hider RC, Cornelis P, Cámara M, Williams P. The *Pseudomonas aeruginosa* 4-quinolone signal molecules HHQ and PQS play multifunctional roles in quorum sensing and iron entrapment. Chem Biol. 2007; 14:87-96.

3. Higgins S, Heeb S, Rampioni G, Fletcher MP, Williams P, Cámara M. Differential Regulation of the Phenazine Biosynthetic Operons by Quorum Sensing in *Pseudomonas aeruginosa* PAO1-N. Front Cell Infect Microbiol. 2018; 8:252.

4. Kulasekara HD, Ventre I, Kulasekara BR, Lazdunski A, Filloux A, Lory S. A novel two-component system controls the expression of *Pseudomonas aeruginosa* fimbrial cup genes. Mol Microbiol. 2005; 55:368-80.

5. Schnider-Keel U, Lejbølle KB, Baehler E, Haas D, Keel C. The sigma factor AlgU (AlgT) controls exopolysaccharide production and tolerance towards desiccation and osmotic stress in the biocontrol agent *Pseudomonas fluorescens* CHA0. Appl Environ Microbiol. 2001; 67:5683-93.

6. Kovach ME, Elzer PH, Hill DS, Robertson GT, Farris MA, Roop RM 2nd, Peterson KM. Four new derivatives of the broad-host-range cloning vector pBBR1MCS, carrying different antibiotic-resistance cassettes. Gene. 1995; 166:175–176

7. Niewerth H, Bergander K, Chhabra SR, Williams P, Fetzner S. Synthesis and biotransformation of 2-alkyl-4(1H)-quinolones by recombinant *Pseudomonas putida* KT2440. Appl Microbiol Biotechnol. 2011; 91:1399-408.

8. Ilangovan A, Fletcher M, Rampioni G, Pustelny C, Rumbaugh K, Heeb S, Cámara M, Truman A, Chhabra SR, Emsley J, Williams P. Structural basis for native agonist and synthetic inhibitor recognition by the *Pseudomonas aeruginosa* quorum sensing regulator PqsR (MvfR). PLoS Pathog. 2013; 9:e1003508.

9. West SE, Schweizer HP, Dall C, Sample AK, Runyen-Janecky LJ. Construction of improved *Escherichia-Pseudomonas* shuttle vectors derived from pUC18/19 and sequence of the region required for their replication in *Pseudomonas aeruginosa*. Gene. 1994; 148:81-6.

10. Rampioni G, Pustelny C, Fletcher MP, Wright VJ, Bruce M, Rumbaugh KP, Heeb S, Cámara M, Williams P. Transcriptomic analysis reveals a global alkyl-quinolone-independent regulatory role for PqsE in facilitating the environmental adaptation of *Pseudomonas aeruginosa* to plant and animal hosts. Environ Microbiol. 2010; 12:1659-73

11. Becher A, Schweizer HP. Integration proficient *Pseudomonas aeruginosa* vectors for isolation of single copy chromosomal *lacZ* and *lux* gene fusions. Biotechniques. 2000; 29:948-50.

12. Figurski D. H., Helinski D. R. Replication of an origin-containing derivative of plasmid RK2 dependent on a plasmid function provided in trans. *Proc Natl Acad Sci U S A.* **76,** 1648-52 (1979).

13. Kaniga K., Delor I., Cornelis G. R. A wide-host-range suicide vector for improving reverse genetics in gram-negative bacteria: inactivation of the *blaA* gene of *Yersinia enterocolitica*. *Gene*. **109,** 137-41 (1991).

14. Hoang T. T., Kutchma A. J., Becher A., Schweizer H. P. Integration-proficient plasmids for *Pseudomonas aeruginosa*: site-specific integration and use for engineering of reporter and expression strains. *Plasmid*. **43,** 59-72 (2000)

**Table S2** Oligonucleotide primers used in this study

| Name | Sequence (5′-3′, restriction sites underlined) |
|---|---|
| TnM1 | GTGAGCGGATAACAATTTCACACAG |
| TnM2 | ACAGGAAACAGGACTCTAGAGG |
| TnMseq | CACCCAGCTTTCTTGTACAC |
| F*tatA* | GGAATTCCCCTGAACCTACACATTGCCA |
| R*tatA* | GGGGTACCCCATTCCGAACATCGATGGCTA |
| Tat3D-UF | TATGAATTCAATCTATTGGTCGCGTTC |
| Tat3D-UR | TATGGATCCAAAAATGCCCATGTCGTA |
| Tat3D-DF | TATGGATCCACCCGCCAGTGAACCTGC |
| Tat3D-DR | TATTCTAGACGCGCAGCTTGTCGACCT |
| S.PA4431UF | CAGGTCGACGGATCCCCGGGGGTACTGGCAAGATTTCCAT |
| S.PA4431DF | TATGCATCCGCGGGCCCGGGGGAGATCAGGAAGTCACCGC |
| S.PA4431DF | GACTGAATGTGATCGGCGTGGACCAGGAG |
| S.PA4431UR | CGCCGATCACATTCAGTCGTCTCCCATCA |
| S-4431CTXFor | TCCCCCGGGCTGCAGGAATTCTTCGACGCTTGCTGAAAA |
| S-4431CTXRev | GATAAGCTTGATATCGAATTCTCAGGCTTTCTCCTGGTC |
| *rhlA* CTX UF | TATAAGCTTTGCCAAAAGCCTGAC |
| *rhlA* CTX DR | TATGGATCCTTGCAAACCGATACC |
| *pqsR* CTX UF | TCCAGCGAATTCGATACGCAACCGCCG |
| *pqsR* CTX DR | GATGACCTGCAGGAACATGTTCACGTG |
| *tac* CTX UF | AAACTCCTCGAGCATCAAATGAAACTG |
| *tac* CTX DR | GAGCTCGAATTCTGTTTCCTGTGTGAA |

**Table S3**. *P. aeruginosa* PA14 Tat substrate mutants used in this study

| Mutant Number | PA Number/Name | Function |
|---|---|---|
| 1 | PA0144 | Nucleoside 2-deoxyribosyltransferase |
| 2 | PA0365 | Hypothetical protein |
| 3 | PA0735 | Hypothetical protein |
| 4 | PA0844 | Hemolytic phospholipase C (PlcH) |
| 5 | PA1174 | Nitrate reductase catalytic subunit (NapA) |
| 6 | PA1601 | Aldehyde dehydrogenase |
| 7 | PA1880 | Oxidoreductase |
| 8 | PA2065 | Copper resistance protein (CopA) |
| 9 | PA2124 | Dehydrogenase |
| 10 | PA2264 | Hypothetical protein |
| 11 | PA2328 | Hypothetical protein |
| 12 | PA2378 | Aldehyde dehydrogenase |
| 13 | PA2389 | Hypothetical protein (PvdR) |
| 14 | PA2392 | Tyrosinase (PvdP) |
| 15 | PA2394 | Aminotransferase (PvdN) |
| 16 | PA2531 | Aminotransferase |
| 17 | PA2635 | Hypothetical protein |
| 18 | PA2699 | Hydrolase |
| 19 | PA3222 | Permease |
| 20 | PA3319 | Non-hemolytic phospholipase C (PlcN) |
| 21 | PA3392 | Nitrous-oxide reductase (NosZ) |
| 22 | PA3713 | Spermidine dehydrogenase (SpdH) |
| 23 | PA3768 | Metallo-oxidoreductase |
| 24 | PA3910 | Phosphodiesterase/alkaline phosphatase (EddA) |
| 25 | PA4140 | Cholesterol oxidase (ChoA) |
| 26 | PA4159 | Iron-enterobactin transporter periplasmic binding protein (FepB) |
| 27 | PA4431 | Cytochrome $bc_1$ Rieske subunit (PetA) |
| 28 | PA4621 | Oxidoreductase |
| 29 | PA4692 | Sulphite oxidase subunit (YedY) |
| 30 | PA4812 | Formate dehydrogenase-O, major subunit (FdnG) |
| 31 | PA4858 | Hypothetical protein |
| 32 | PA5327 | Oxidoreductase (SphC) |
| 33 | PA5538 | N-acetylmuramoyl-L-alanine amidase AmiC* |
| 34 | PA14_48450 | Peptidyl-arginine deiminase (Agu2A')** |

Annotations are based on predictions from the website www.pseudomonas.com or are inferred from sequence homology. Functional predictions by Gimenez *et al*. (2018) Scientific Reports 11950. doi: 10.1038/s41598-018-30393-x.
*Annotated as AmiA on *Pseudomonas* PA14 genome but sequence is closer to that of AmiC.
** No conserved gene found in PAO1 so retains the PA14 annotation.

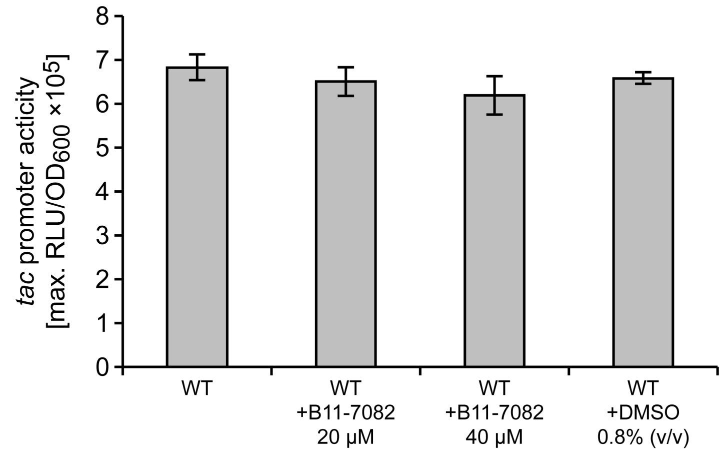


**Fig. S1** The Tat inhibitor Bayer 11-7082 has no effect on light output at 20 or 40 µM from *P. aeruginosa* CTX:*ptac′-luxCDABE* chromosomal reporter fusion. The co-solvent DMSO, had no effect at 0.8% on the *lux* reporter fusion either. Data are presented as maximal light output as a function of growth (RLU/$OD_{600}$). Experiments were repeated in triplicate at least twice.

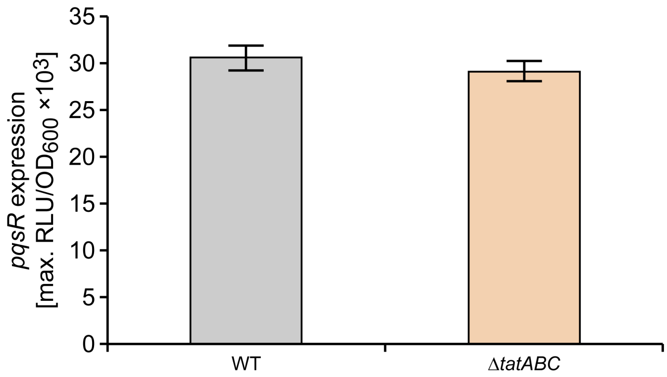


**Fig**. **S2** Deletion of the *P. aeruginosa tatABC* genes does not influence *pqsR* expression. The data show that there are no differences in the expression of a chromosomal CTX::*pqsR′-luxCDABE* fusion in the *P. aeruginosa* wild type compared with the Δ*tatABC* mutant Data are presented as maximal light output as a function of growth (RLU/$OD_{600}$). Experiments were repeated in triplicate at least twice.

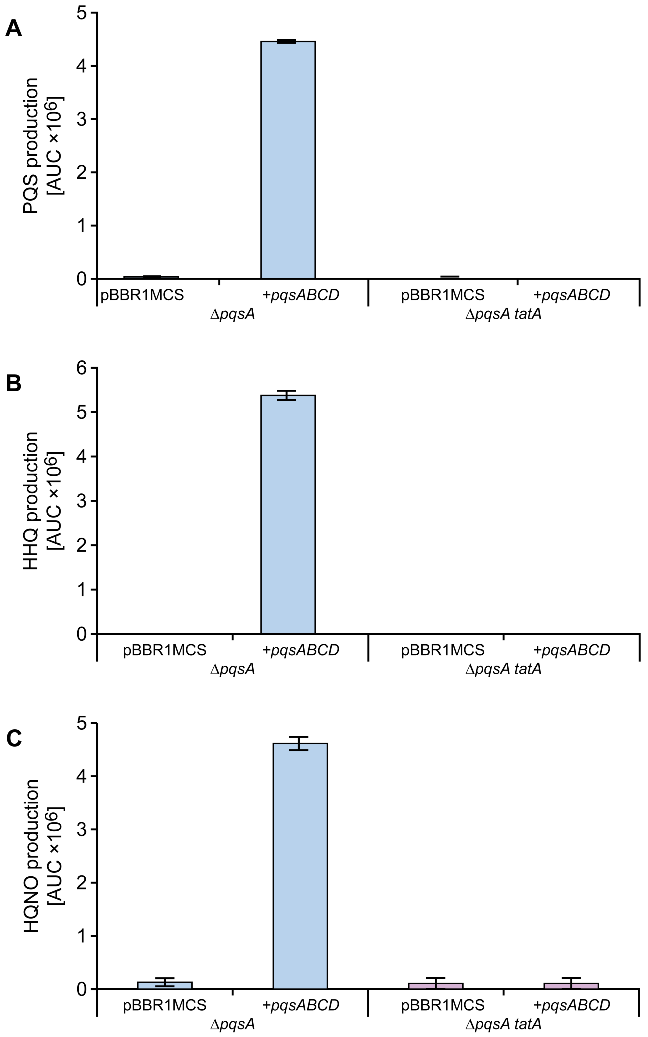


**Fig.S3** AQ biosynthesis is not restored in a *tatA* Δ*pqsA* double mutant by plasmid-borne *pqsABCD* in the absence of autoinduction. Semi-quantitative analysis by LC-MS/MS of PQS, HHQ and HQNO production by *P. aeruginosa pqsA* and *tatA* Δ*pqsA* mutants respectively without or with the *pqsABCD* biosynthetic genes provided *in trans* via pBBRMCS::*pqsABCD.* Experiments were repeated in triplicate at least twice.

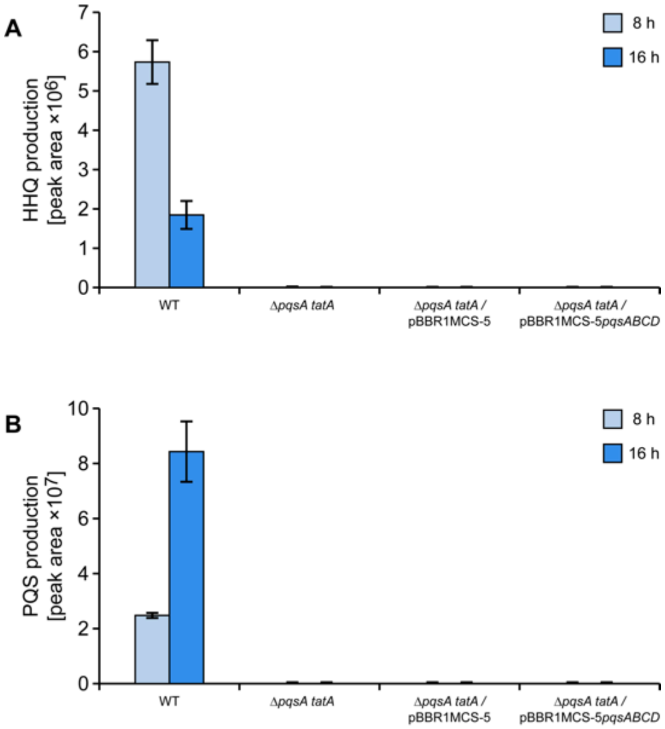


**Fig. S4.** HHQ and PQS do not accumulate intracellularly a *tatA* Δ*pqsA* double mutant harboring the plasmid-borne *pqsABCD* genes in the absence of autoinduction*.* Semi-quantitative analysis by LC-MS/MS of PQS (**A**) and HHQ (**B**) extracted from whole cells of *P. aeruginosa* wild type and the *tatA* Δ*pqsA* mutant without or with the *pqsABCD* biosynthetic genes provided via pBBRMCS::*pqsABCD.* Cells were harvested at 8 h and 16 h respectively*.* Experiments were repeated in triplicate.

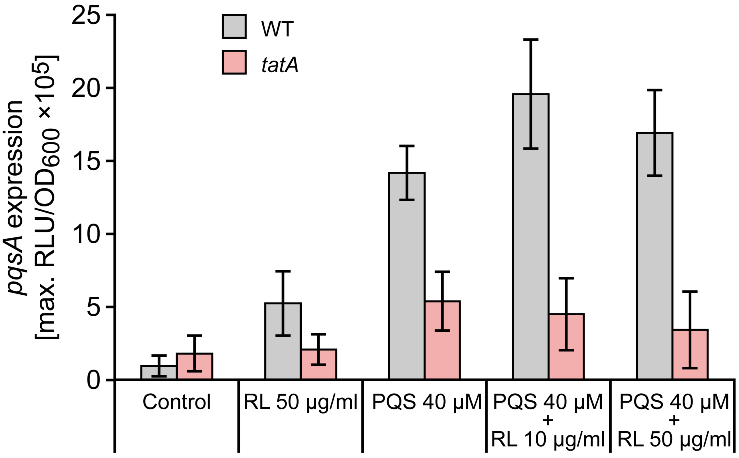


**Fig. S5.** Exogenous rhamnolipids do not enhance PQS-dependent expression of *pqsA* in a *tatA* Δ*pqsA* mutant*.* PQS (40 µM) was added with or without purified rhamnolipids (50 µg/ml) to a *pqsA* mutant or a *tatA* Δ*pqsA* mutant carrying chromosomal *pqsA′-lux* fusions. Maximal light output as a function of growth (RLU/$OD_{600}$) is presented. Experiments were repeated in triplicate at least twice.

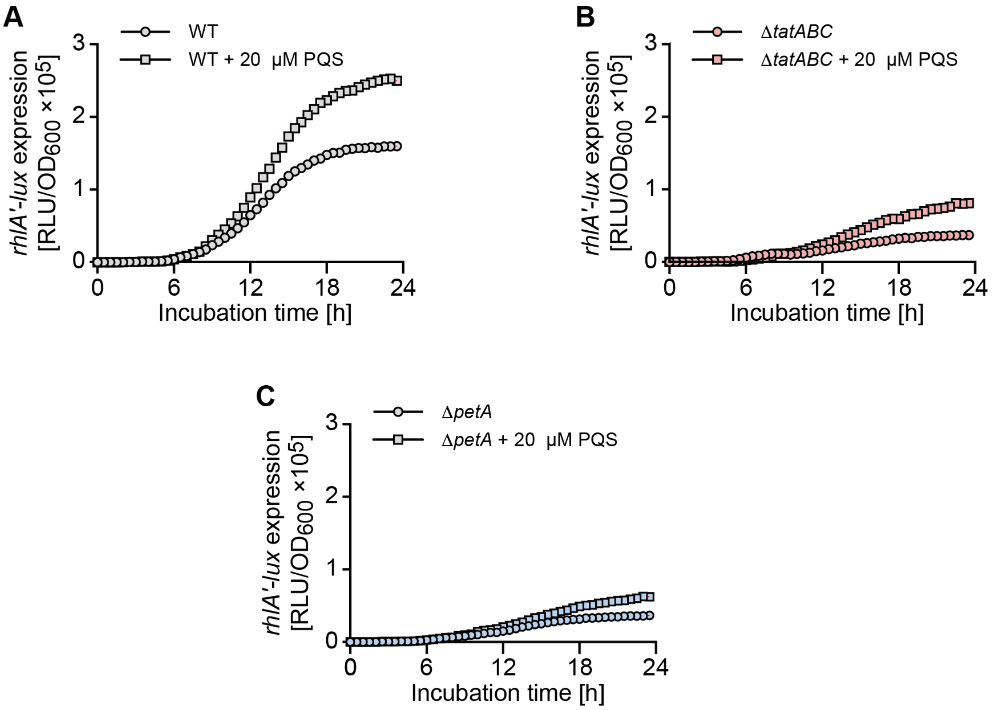


**Fig. S6.** Rhamnolipid biosynthesis gene *rhlA* shows altered expression profiles in *P. aeruginosa* PA14 Δ*tatABC* and Δ*petA* mutants compared with wild type and fail to respond to exogenous PQS. Bioluminescence from a chromosomal *rhlA′-lux* fusion as a function of growth (RLU/OD) over time when introduced into (**A**) the PA14 wild type, (**B**) Δ*tatABC* and (**C**) Δ*petA* mutants in the absence or presence of exogenous PQS (20 μM).

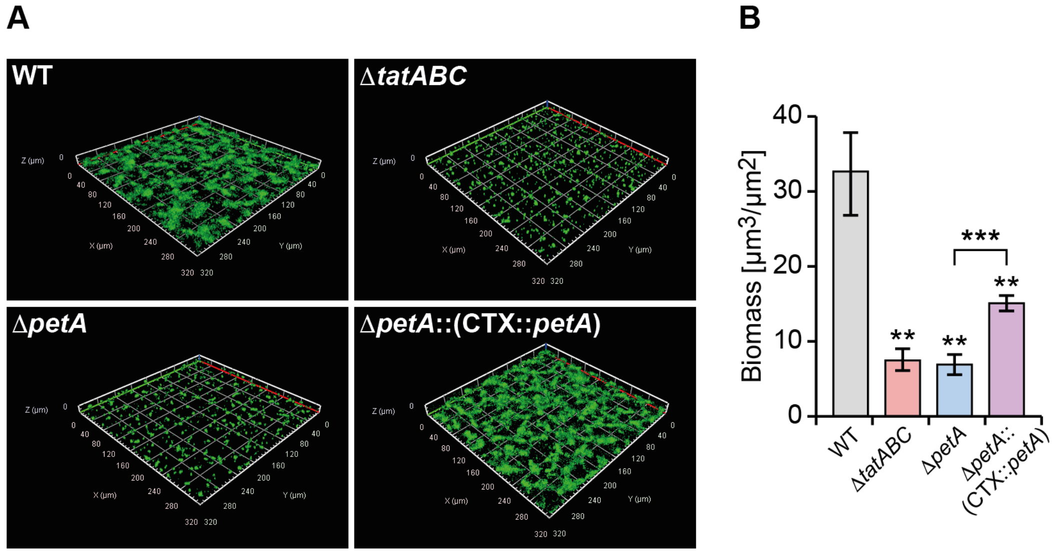


**Fig. S7.** Deletion of *petA* in *P. aeruginosa* PA14 results in a reduction in the eDNA content of biofilms. Biofilms of wild type PA14, the Δ*tatABC* and Δ*petA* mutants and the genetically complemented PA14 mutant (Δ*petA*::(CTX::*petA*)) were grown cultured statically and stained for eDNA with YOYO-1. (**A**) CLSM images and (**B**) eDNA quantification. Experiments were repeated in triplicate at least twice. ***$p < 0.001$, **$p < 0.01$.
